# The CD58:CD2 axis is co-regulated with PD-L1 via CMTM6 and governs anti-tumor immunity

**DOI:** 10.1101/2022.03.21.485049

**Authors:** Patricia Ho, Johannes C. Melms, Meri Rogava, Chris J. Frangieh, Shivem B. Shah, Zachary Walsh, Oleksandr Kyrysyuk, Amit Dipak Amin, Lindsay Caprio, Benjamin T. Fullerton, Rajesh Soni, Casey R. Ager, Jana Biermann, Yiping Wang, Michael Mu, Hijab Fatima, Emily K. Moore, Neil Vasan, Samuel F. Bakhoum, Steven L. Reiner, Chantale Bernatchez, Emily M. Mace, Kai W. Wucherpfennig, Dirk Schadendorf, Gary K. Schwartz, Benjamin Izar

**Author notes:** contributed equally to this work (Patricia Ho, Johannes C. Melms, Meri Rogava).

## Abstract

The cell autonomous balance of immune-inhibitory and -stimulatory signals is a critical yet poorly understood process in cancer immune evasion. Using patient-derived co-culture models and humanized mouse models, we show that an intact CD58:CD2 interaction is necessary for anti-tumor immunity. Defects in this axis lead to multi-faceted immune evasion through impaired CD2-dependent T cell polyfunctionality, T cell exclusion, impaired intra-tumoral proliferation, and concurrent protein stabilization of PD-L1. We performed genome-scale CRISPR-Cas9 and CD58 coimmunoprecipitation mass spectrometry screens identifying CMTM6 as a key stabilizer of CD58, and show that CMTM6 is required for concurrent upregulation of PD-L1 in *CD58* loss. Single-cell RNA-seq analysis of patient melanoma samples demonstrates that most TILs lack expression of primary costimulatory signals required for response to PD-1 blockade (e.g. *CD28*), but maintain strong *CD2* expression, thus providing an opportunity to mobilize a so far therapeutically untapped pool of TILs for anti-tumor immunity. We identify two potential therapeutic avenues, including rescued activation of human *CD2*-expressing TILs using recombinant CD58 protein, and targeted disruption of PD-L1/CMTM6 interactions. Our work identifies an underappreciated yet critical axis at the nexus of cancer immunity and evasion, uncovers a fundamental mechanism of co-inhibitory and -stimulatory signal balancing, and provides new approaches to improving cancer immunotherapies.

## INTRODUCTION

Immune checkpoint blockade (ICB) therapies, such as anti-PD-1 antibodies, enable CD8+ T cell-mediated anti-tumor immunity and produce durable responses in a portion of patients with metastatic melanoma and other cancers^1,2^; however, most patients are either intrinsically resistant or acquire resistance after an initial response. While there are a few clinically validated mechanisms of ICB resistance, including genomic alterations that result in impaired antigen presentation or mutations in the IFN-ɣ-JAK/STAT pathway^3-5^, no clear mechanism of cancer immune evasion is identified in most patients.

Using single-cell RNA-seq in melanoma patients, we recently found that downregulation of *CD58* (also known as lymphocyte adhesion factor 3, *LFA-3*) is associated with resistance to ICB and inferred T cell exclusion^6-8^; however, the underlying mechanisms are poorly understood. CD58 is physiologically expressed on antigen presenting cells, namely macrophages, where it ligates with its T cell receptor CD2 to deliver “Signal 2” following T cell receptor (TCR) and MHC-presented antigen engagement^9,10^. CD58 further functions as an adhesion molecule, facilitating the initial binding of effector T cells^11-13^. CD58 exists in two isoforms via RNA splicing, a glycosylphosphatidylinositol (GPI)-anchored form and a transmembrane (TM) form (hereto referred to as CD58-GPI and CD58-TM, respectively)^14,15^, though a functional difference between the two isoforms is not well established. The role of CD58 in cancer remains poorly understood, in part due to the lack of a known mouse homolog of *CD58*. Thus, murine models frequently used to study tumor-immune interactions and ICB responses have limited utility in studying the CD58:CD2 axis.

Notably, T cell expression of the primary co-stimulatory protein CD28 is required for response to PD-1 blockade, but CD28 is frequently absent from tumor-infiltrating CD8+ T lymphocytes (TILs)^16-18^. Importantly, in the context of *CD28* loss in human CD8+ T cells, the most potent alternative costimulatory axis for T cell cytokine production and cytotoxicity is CD58 ligation with CD2^19^. As we show here, most TILs lack expression of *CD28*, and therefore do not contribute as a substrate for response to PD-1 blockade; however, they do maintain *CD2* expression. Thus, in principle, engagement of CD28-CD8+ cells via CD58:CD2 ligation may induce T cell activation and overcome resistance to PD-1 blockade.

Here, we show that intact cancer cell *CD58* expression is required for effective T cell mediated antitumor responses, and that loss of *CD58* confers cancer immune evasion through multiple mechanisms, including impaired activation and proliferation of T cells, reduced tumor lytic capacity, reduced tumor infiltration and intratumoral T cell expansion, and concurrent upregulation of coinhibitory PD-L1. To determine the mechanisms of *CD58* regulation and its coregulation of PD-L1, we performed genome-scale CRISPR-Cas9 and proteomics screens and identify CMTM6 as a novel and critical regulator of CD58 protein abundance. We show that CMTM6 is required for CD58 maintenance by regulating endolysosomal protein recycling, and necessary for concurrently increased PD-L1 in *CD58* loss. We further identify the N-terminal domain of PD-L1 as a binding site for CMTM6 and show that disruption of this interaction results in PD-L1 degradation. Therapeutic stimulation of the CD58:CD2 axis in TILs confers improved anti-tumor immunity. Thus, our work identifies a critical and understudied axis in cancer immunity and reveals a mechanistic balance between co-inhibitory and co-stimulatory signals in the cancer immune synapse. Selective targeting of the CD58:CD2 axis may represent a rational therapeutic approach for mobilization of an untapped pool of TILs in cancer immunotherapy.

## RESULTS

### Intact cancer cell *CD58* expression is necessary for anti-tumor immunity

Prior reports indicated that *CD58* downregulation or loss is associated with cancer immune evasion and resistance to ICB in melanoma or CAR-T cell therapy in lymphoma^6-8,20^; however, the underlying mechanisms are poorly understood. Analysis of The Cancer Genome Atlas (TCGA) indicates that baseline *CD58* expression is variable in cutaneous melanoma and virtually absent in uveal melanoma, a rare subtype of melanoma that arises in the eye and has an extremely low response rate to immunotherapies (Fig. S1A), supporting the notion that cancer immune evasion is associated with low *CD58* expression. In a renal cell carcinoma (RCC) single-cell data set, immune checkpoint exposure was associated with reduced *CD58* expression (Fig. S1B)^21^. Analysis of single-cell RNA-seq data of melanoma patient samples^6,22^ reveals that virtually all TILs maintain high *CD2* expression independent of their differentiation state, suggesting that defects in the CD58:CD2 axis are driven mainly by *CD58* loss or downregulation (Fig. S1C-D).

CD28 is the primary co-stimulatory protein in humans and is required for response to PD-1 blockade^16,17^. In the absence of CD28 on CD8+ T cells, the CD58:CD2 axis is the most potent co-stimulatory pathway to activate T cells^19^. We find that most CD8+ TILs in melanoma lack *CD28* mRNA expression, including progenitor-like and differentiated T cells (Fig. S1C-D), but that *CD28* expression is enriched in responders compared to non-responders to anti-PD-1 therapy within the *TCF7*+ progenitor-like T cell population, per prior reports. Furthermore, we find that *CD28-CD8+* T cells exist within the *TCF7*+ T cell population, suggesting that a large pool of TILs exist that may be amenable to activation via preserved *CD2* expression. In RCC, we also find that expression of *CD2* is preserved in virtually all TILs, while *CD28* expression is lost (Fig. S1E).

To mechanistically dissect the role of *CD58* cancer cell expression in T cell mediated tumor lysis, we used patient-derived co-cultures composed of melanoma cells and autologous, *ex vivo* expanded TILs. We generated otherwise isogenic cell lines with *CD58* knockout (KO) followed by rescue with overexpression (OE) of either *CD58*-TM or *CD58*-GPI open reading frames (ORFs) (Fig. S2A); these lines were then co-cultured with autologous TILs. *CD58* loss resulted in significantly reduced tumor lysis and IFN-ɣ production by T cells, while rescue with either *CD58*-TM or *CD58*-GPI re-sensitized cancer cells to T cell-mediated tumor killing and rescued IFN-ɣ production (Fig. 1A-B, Fig. S2B).

**Figure 1:**
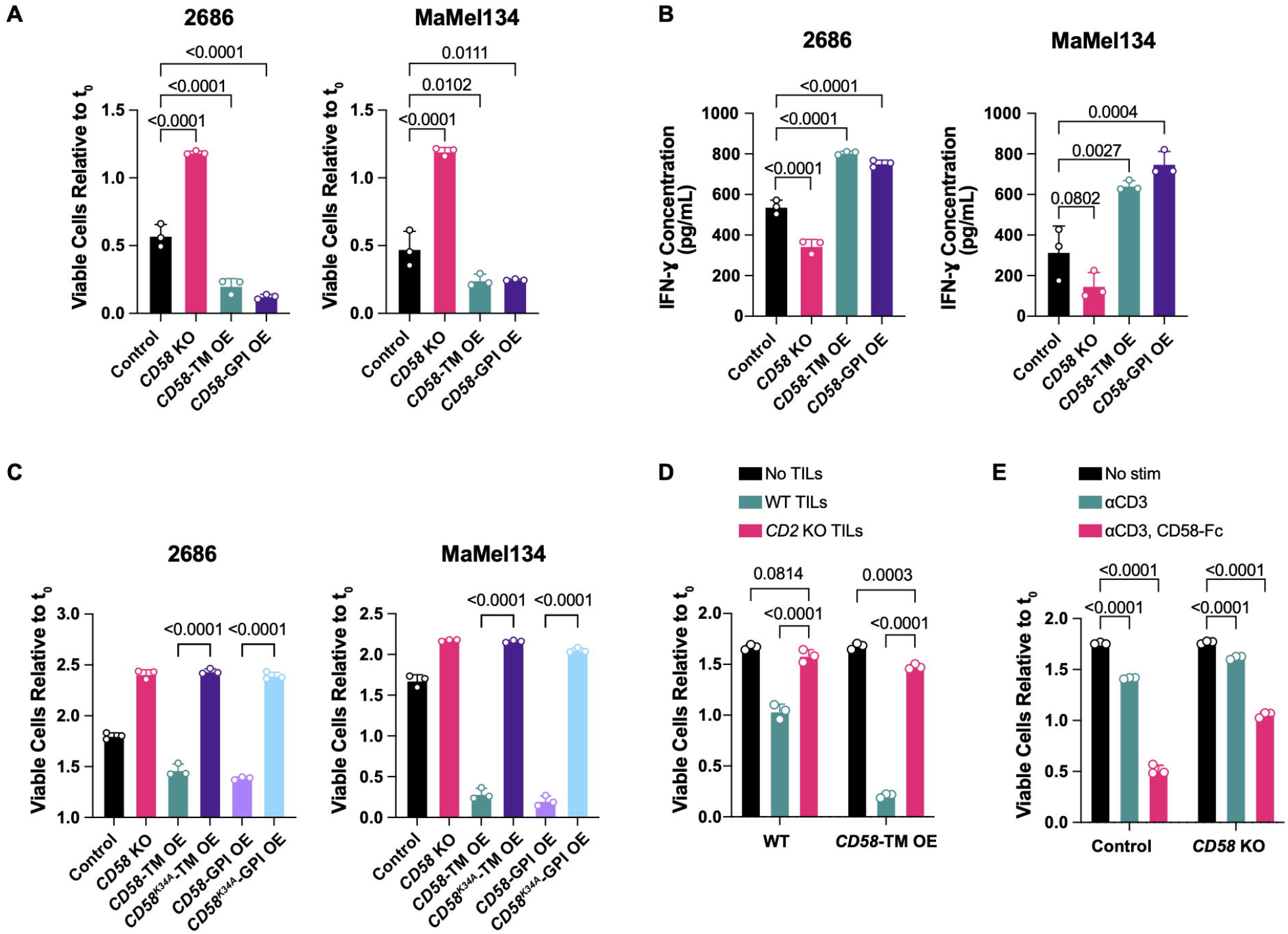
Intact cancer cell CD58 and T cell CD2 signaling is required for anti-tumor immunity. (A-B) Fold change in number of viable 2686 and MaMel134 control, *CD58* KO, and *CD58-*TM or *CD58-*GPI OE cells after 48 h or 72 h co-culture with autologous TILs, respectively (A), and IFN-ɣ concentration within cleared media collected from co-cultures (B). (C) Fold change in number of viable 2686 and MaMel134 control, *CD58* KO, *CD58-*TM OE, *CD58*^*K34A*^-TM OE, *CD58*-GPI OE, or *CD58*^*K34A*^-GPI OE cells after 48 h or 72 h co-culture, respectively, with engineered NY-ESO-1-specific T cells or autologous TILs, respectively. (D) Fold change in number of viable MaMel134 control or *CD58*-TM OE cells after 48 h co-culture with or without WT or *CD2* KO autologous TILs. (E) MaMel134 TILs were stimulated for 48 h with 1 µg/mL OKT3 +/-2 µg/mL CD58-Fc chimera prior to co-culturing with autologous melanoma cells; fold change in viable melanoma cells shown after 48 h of co-culture. Experiments performed in triplicate, with representative experiment shown of at least two independent experiments each. Statistical analysis performed using one-way ANOVA with Tukey’s multiple comparisons test (A-C), and two-way ANOVA with Tukey’s multiple comparisons test (D,E). Data represent mean ± SD.

To exclude the possibility that *CD58* loss confers immune evasion through non-specific effects, we CRISPR-engineered human peripheral blood mononuclear cells to express a TCR with high affinity against the common cancer testis antigen NY-ESO-1 (Fig. S2C)^23^, which is frequently expressed and presented in melanomas with the HLA-A*2:01 allele, including in the cell line 2686. In this engineered co-culture system, loss of either *CD58* or *B2M* (which is required for MHC Class I antigen presentation) in 2686 cells conferred resistance to NY-ESO-1-TCR T cells (Fig. S2D), thus confirming that the *CD58* phenotype was mediated by a specific TCR/epitope interaction, rather than non-specific mechanisms. Importantly, this demonstrates that antigen presentation was not sufficient to promote anti-tumor immunity in the context of *CD58* loss. Together, these results show that *CD58* expression on cancer cells determines tumor sensitivity by modulating cytotoxic TIL activity in a TCR-specific/epitope-dependent manner.

### Cancer cell intrinsic expression of *CD58* modulates T cell responses in a CD2-dependent manner

We next sought to establish that immune modulation by cancer cell autonomous *CD58* expression occurs specifically through interaction with CD2 on T cells. While re-expression of *CD58*-TM or *CD58*-GPI rescues sensitivity of *CD58* KO cells to co-culture with T cells, addition of either a CD2- or CD58-blocking antibody (but not IgG control) abrogates this sensitization (Fig. S2E-F). Orthogonally, we used site-directed mutagenesis to create K34A mutated *CD58*^K34A^-TM and *CD58*^K34A^-GPI ORFs, which show impaired binding to CD2 (Fig. S2G-H)^24^. We rescued *CD58* KO cells with *CD58*^WT^-TM, *CD58*^WT^-GPI, *CD58*^K34A^-TM, or *CD58*^K34A^-GPI ORFs, followed by co-culture with autologous TILs or NY-ESO-1-TCR T cells. While overexpression of WT *CD58* constructs rescued sensitivity to killing by TILs and IFN-ɣ production, overexpression of K34A mutant *CD58* did not (Fig. 1C, Fig. S2I). Next, we used CRISPR-Cas9 to knockout *CD2* on TILs (Fig. S2J), and co-cultured WT or *CD2* KO TILs with parental or *CD58*-TM OE cancer cells. We find that, compared to WT TILs, *CD2* KO TILs demonstrated minimal cytotoxicity against either parental or *CD58*-TM cells (Fig. 1D).

Next, we considered whether TILs can be stimulated via CD2 to enhance their cytotoxic activity. We utilized a recombinant CD58-Fc chimera homodimer, which in combination with CD3 stimulation was sufficient to increase TIL proliferation and IL-2 production *in vitro*, more so than CD3 stimulation alone or in combination with CD28 stimulation (Fig. S2K-L). Notably, cells receiving co-stimulation from the CD58-Fc chimera (compared to CD3 activated TILs) more efficiently killed parental cancer cells (as well as *CD58* KO cells to a lesser extent, likely due to bystander cytokine activity) (Fig. 1E). Lastly, there was no expression of CD48, a previously proposed alternative CD2 ligand, on multiple melanoma cell lines; thus, CD48 is unlikely to play an important role in this context (Fig. S1M). Together, these results show that an intact CD58:CD2 axis is necessary for efficient TIL-mediated lysis of cancer cells, and that disruption of this specific ligand or receptor interaction confers cancer immune evasion.

### Loss of *CD58* confers cancer immune evasion via impaired intratumoral T cell infiltration and proliferation

A major mechanism of resistance to ICB is poor tumor infiltration of T cells or exclusion of T cells from the tumor-microenvironment (TME)^25,26^. We therefore sought to examine whether *CD58* loss could also impair T cell infiltration of tumors *in vivo*. Because there is no known mouse homolog of *CD58*, we established a partly humanized mouse model using the severely immunodeficient NOG mouse (NOD.Cg-*Prkdc*^*scid*^ *Il2rg*^*tm1Sug*^/JicTac) with transgenic expression of human interleukin 2 (h*IL2*), which is required for T cell survival (Fig. S3A). In these animals, we subcutaneously implanted *CD58* WT melanoma cells alongside *CD58* KO or *CD58*-TM OE cells (n=16 each) to form bilateral flank tumors followed by adoptive cell transfer (ACT) with autologous TILs or phosphate buffered saline (PBS) control (Fig 2A). *CD58* KO tumors were highly resistant to ACT (with no responders among eight animals), while half of *CD58* WT tumors demonstrated either a complete or partial response to ACT (Fig. 2B-C), suggesting that *CD58* loss drives immune evasion *in vivo*.

**Figure 2:**
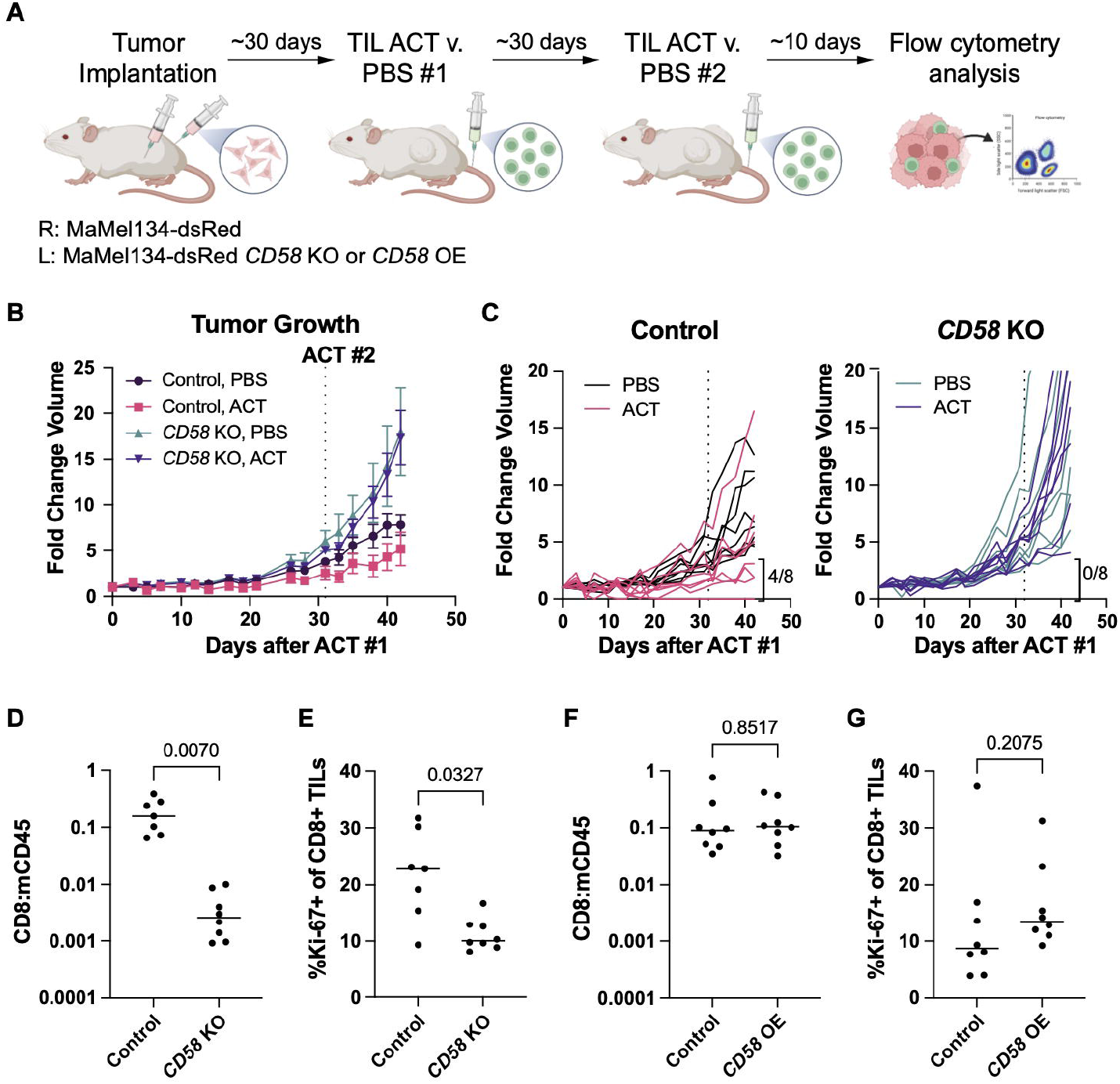
Loss of *CD58* confers cancer immune evasion via impaired intratumoral T cell infiltration and proliferation. (A) Experimental design of *in vivo* study of *CD58* loss and re-expression in melanoma tumors. MaMel134 NLS-dsRed-expressing parental and *CD58*-KO or *CD58*-TM OE melanoma cells were implanted in NOG mice as bilateral subcutaneous flank injections, followed by two treatments with adoptive cell transfer of autologous TILs or PBS control (n=8 for each treatment group). Tumors were collected and processed for flow cytometry analysis of tumor infiltration by TILs. Created in BioRender. (B) Fold change growth of control and *CD58* KO tumors following initial ACT treatment or PBS control. (C) Individual fold change growth of control and *CD58* KO tumors following initial ACT treatment or PBS control. For each group, individual tumors with partial or complete response to therapy, defined as <4-fold change in volume from initial ACT treatment to endpoint, are indicated for each group, (D) Ratio of human CD8+ cells compared to number of mouse CD45+ cells, in order to account for tumor size, within ACT-treated control and *CD58* KO tumors, as analyzed by flow cytometry. (E) Percent of CD8+ TILs within ACT-treated control and *CD58* KO tumors that express Ki-67, as analyzed by flow cytometry. (F) As in (D), but comparing control v. *CD58* OE tumors. (G) As in (E), but comparing control v. *CD58* OE tumors. Statistical analysis performed using paired two-sided T-tests. Line at median (D-G). Data represent mean ± SEM (B).

We next analyzed tumors by flow cytometry for T cell infiltration and phenotypes (Fig. S3B) and found that nearly all surviving TILs within tumors were CD8+ (Fig. S3C-E). *CD58* KO tumors showed ∼100 fold lower infiltration by CD8+ TILs compared to *CD58* WT tumors (Fig. 2D). We further established that reduced TIL abundance in *CD58* KO tumors was not simply a function of larger tumor size, as infiltration was minimal across a spectrum of tumor sizes compared to *CD58* WT tumors (Fig. S3F). Furthermore, among tumor-infiltrating T cells, those from *CD58* KO tumors showed a significantly reduced proliferation rate compared to *CD58* WT tumors (Fig. 2E). Importantly, rescue of *CD58* KO tumors with *CD58* OE reversed these observations and resulted in intratumoral T cell infiltration and proliferation similar to that found in *CD58* WT tumors (Fig. 2F-G). Furthermore, *CD58* OE tumors had a greater percentage of CD28-CD8+ TILs than *CD58* WT tumors (Fig. S3G), a difference not seen between *CD58* WT and *CD58* KO tumors (Fig. S3H), suggesting that overexpression of *CD58* in these tumors either recruits greater numbers of CD28-CD8+ T cells, or promotes greater T cell activation, leading to *CD28* downregulation.

All together, these results demonstrate that *CD58* expression is necessary for infiltration by cytotoxic T cells in melanoma, and that loss of *CD58* confers resistance to ACT.

### Concurrent upregulation of PD-L1 in *CD58* loss contributes to cancer immune evasion

We previously noted that *CD58* loss was sufficient to promote concurrent upregulation of PD-L1^8^, but the underlying mechanisms and functional role of this co-regulation remain unknown. To address this, we first used *CD58* KO models that were rescued with either *CD58*-TM or *CD58*-GPI, followed by IFN-ɣ stimulation and flow-cytometric or total protein evaluation of PD-L1. While *CD58* loss resulted in increased PD-L1 surface and total protein abundance, rescue with either *CD58* isoform abrogated concurrent PD-L1 increases, even suppressing PD-L1 to below baseline levels (Fig. 3A-B, Fig. S4A-B). Interestingly, *CD58* KO cells rescued with K34A-mutated *CD58* ORFs were also able to downregulate PD-L1 expression (Fig. 3C), suggesting that this CD2-binding site is not necessary for CD58/PD-L1 co-regulation. Importantly, cell surface expression of MHC I protein remained unchanged in both *CD58* null cells and with rescue of either isoform of *CD58* (Fig. 3D, Fig. S4C), indicating that neither resistance nor rescued sensitivity to TIL co-culture (Fig. 1A) were due to changes in expression of antigen presentation genes. These results suggest that expression of PD-L1 is co-regulated with that of CD58.

**Figure 3:**
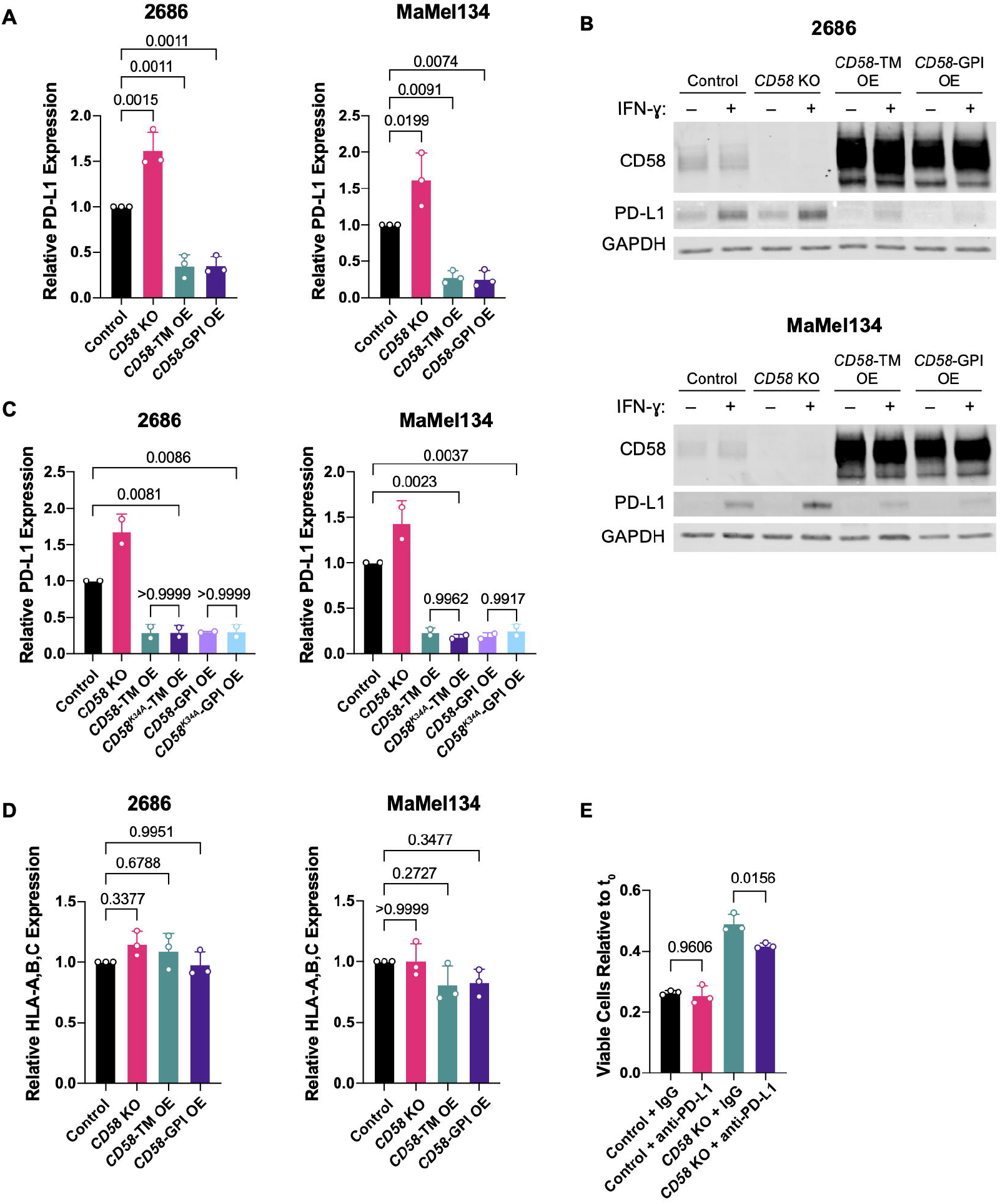
Concurrent upregulation of PD-L1 in *CD58* loss contributes to cancer immune evasion. (A-B) Surface (A) and whole protein (B) PD-L1 expression in 2686 and MaMel134 control, *CD58* KO, and *CD58-*TM or *CD58*-GPI OE cells following 72 h stimulation with 10 ng/mL IFN-ɣ assessed by flow cytometry and immunoblotting, respectively. (C) As in (A), but in 2686 and MaMel134 control, *CD58* KO, *CD58-*TM, *CD58*^*K34A*^-TM, *CD58*-GPI, or *CD58*^*K34A*^-GPI OE cells. (D) As in (A), but showing HLA-A,B,C expression. (E) Fold change in number of viable MaMel134 control and *CD58* KO cells after 24 h co-culture with pre-stimulated autologous TILs in the presence of 10 µg/mL anti-PD-L1 (B7-H1) blocking antibody or IgG isotype control. TILs were stimulated overnight with 1 µg/mL anti-CD3 (OKT3) antibody prior to co-culture with melanoma cells to promote PD-1 expression. Experiments performed in duplicate (A, C, D) or triplicate (E). Independent experiments shown in (A, C, D) and representative experiment shown from at least two independent experiments in (E). Representative blots shown from two independent experiments each (B). Statistical analysis performed using one-way ANOVA with Tukey’s multiple comparisons test. Data represent mean ± SD.

To determine whether increased PD-L1 contributes to cancer immune evasion in *CD58* loss, we next performed co-culture experiments of *CD58* KO cancer cells with or without pharmacological PD-L1 blockade. For this purpose, we first generated “exhausted” CD8+ autologous TILs via chronic TCR stimulation *in vitro* (**Methods**), resulting in PD-1^high^ TILs (Fig. S4D). When co-cultured with exhausted TILs in the presence of an anti-PD-L1 blocking antibody, *CD58* KO melanoma cells demonstrate increased sensitivity to TIL-mediated killing, whereas control cells remain unaffected (Fig. 3E). Thus, elevated PD-L1 expression in *CD58* loss contributes to immune evasion, adding to the several mechanisms through which *CD58* loss or downregulation may contribute to resistance to ICB.

### Genome-scale CRISPR/Cas9 screen identifies *CMTM6* as a regulator of CD58

The regulation of CD58 is poorly understood. To systematically discover regulators of CD58, we performed a genome-wide CRISPR/Cas9 loss-of-function screen combined with fluorescence-activated cell sorting (FACS) (Fig. 4A). We utilized the pooled lentiviral single-guide RNA (sgRNA) library Brunello, which contains 76,441 sgRNAs (4 sgRNAs/gene and 1000 non-targeting control sgRNAs)^27^. A375 melanoma cells with stable expression of Cas9 were transduced with sgRNAs at a multiplicity of infection (MOI) of less than 0.3 for a representation of no more than one sgRNA per cell, with ∼500 cells per sgRNA (Fig. S5A-B). On day 10 post-transduction, cells were stained with anti-CD58-APC antibody, and cells with either low CD58 expression (bottom 5%, CD58^lo^) or within one standard deviation of the mean intensity (CD58^mi^) were collected by FACS (Fig. S5C-D), followed by genomic DNA isolation, sgRNA amplification by PCR, and next generation sequencing (Fig. S5E).

**Figure 4:**
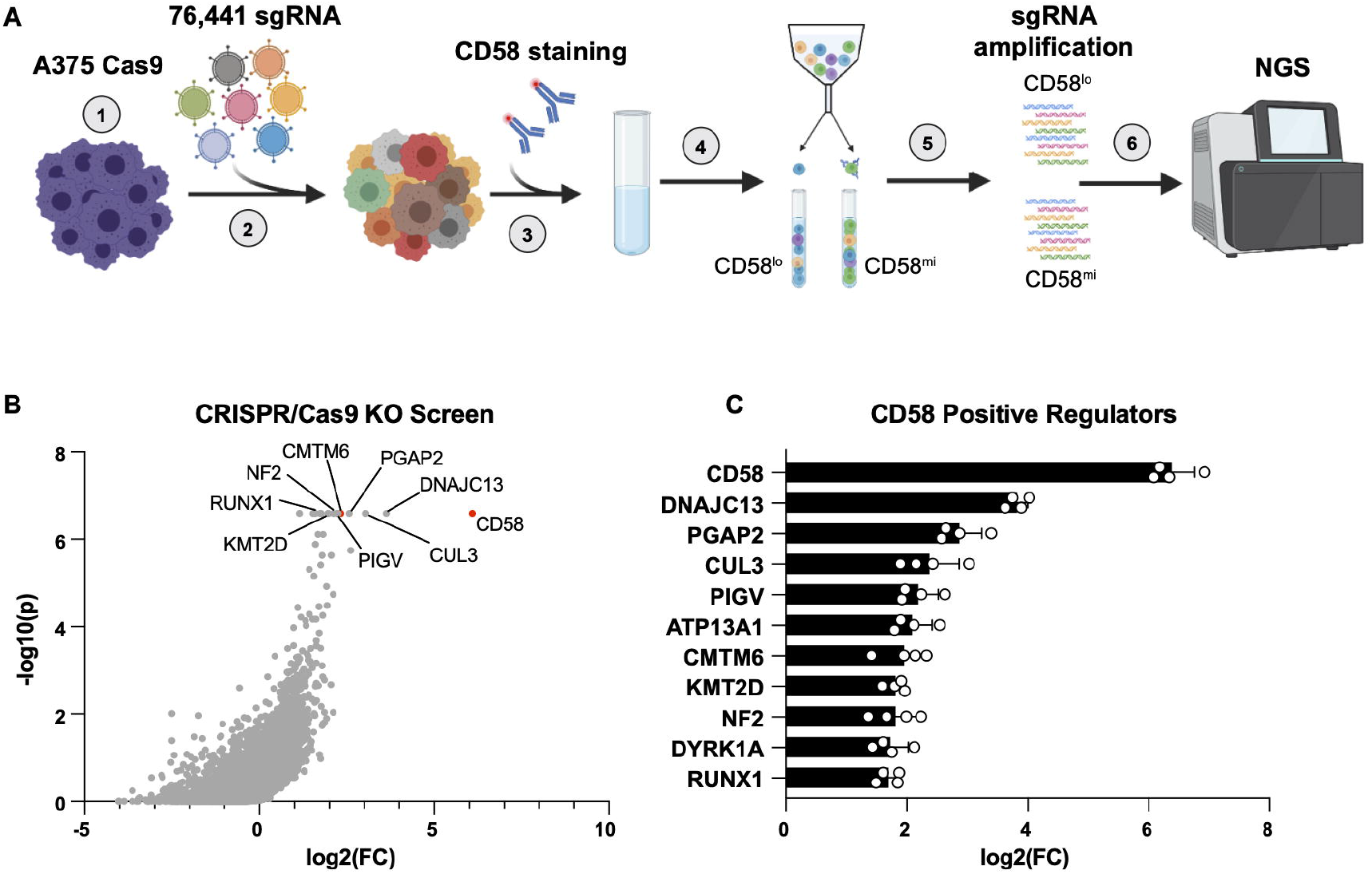
Genome-scale CRISPR/Cas9 screen identifies CMTM6 as regulator of CD58. (A) Experimental design of whole genome-wide CRISPR/Cas9 KO screen to identify positive regulators of *CD58*. 1.) A375 melanoma cells expressing Cas9 were generated and then 2.) transduced with Brunello sgRNA library and cultured for 10 days. 3.) Cells were stained with anti-CD58-APC antibody, and 4.) cells with low surface expression of CD58 (CD58^lo^) were collected alongside a control CD58^mi^ population by FACS, followed by 5.) PCR amplification of integrated sgRNAs and 6.) next generation sequencing (NGS). Screen was performed independently twice, with two technical replicates each. (B) Volcano plot demonstrating distribution of CRISPR/Cas9 target gene enrichment within the CD58^lo^ population compared to the CD58^mi^ population from an example replicate, showing log2 fold change of gene enrichment and P-value for enrichment calculated from a negative binomial model as determined by the MAGeCK algorithm. (C) Genes identified as positive regulators of CD58 (criterion of FDR<0.25 across all four replicates), ranked by log2 fold change enrichment of targeting sgRNAs in CD58^lo^ versus control CD58^mi^ population across replicates. Data represent mean ± SD.

The premise of this screen is that perturbations enriched in the CD58^lo^ population may be physiological regulators of CD58. Enrichment of sgRNAs within the CD58^lo^ population relative to the CD58^mi^ population was computed using MAGeCK (Model-based Analysis of Genome-wide CRISPR/Cas9 Knockout), which identifies enriched sgRNAs, genes, and pathways in CRISPR/Cas9 KO screens^28^ (Fig. 4B, Table S1,2). Eleven gene targets were consistently enriched within CD58^lo^ cells compared to CD58^mi^ cells (Fig. 4C, Fig. S5F). As expected, the top “hit” in this screen was knockout of *CD58* itself; other expected hits included *PIGV* and *PGAP2*, which are both involved in GPI anchor formation, further confirming the robustness of this screen. Interestingly, among the top hits was *CMTM6*, which has recently been described as an important regulator of PD-L1 maintenance^29,30^.

### CMTM6 binds to and promotes protein stability of CD58 via endosomal recycling

To complement this genetic approach, we also performed mass spectrometry analysis of CD58 immunoprecipitation lysates (MS-IP) to identify novel regulators of CD58 on a protein level. Across three biological replicates in two different human melanoma cell lines (A375 and 2686), CMTM6 was consistently identified as a protein interactor with CD58 (Fig. 5A, Fig. S6A-B, Table S3).

**Figure 5:**
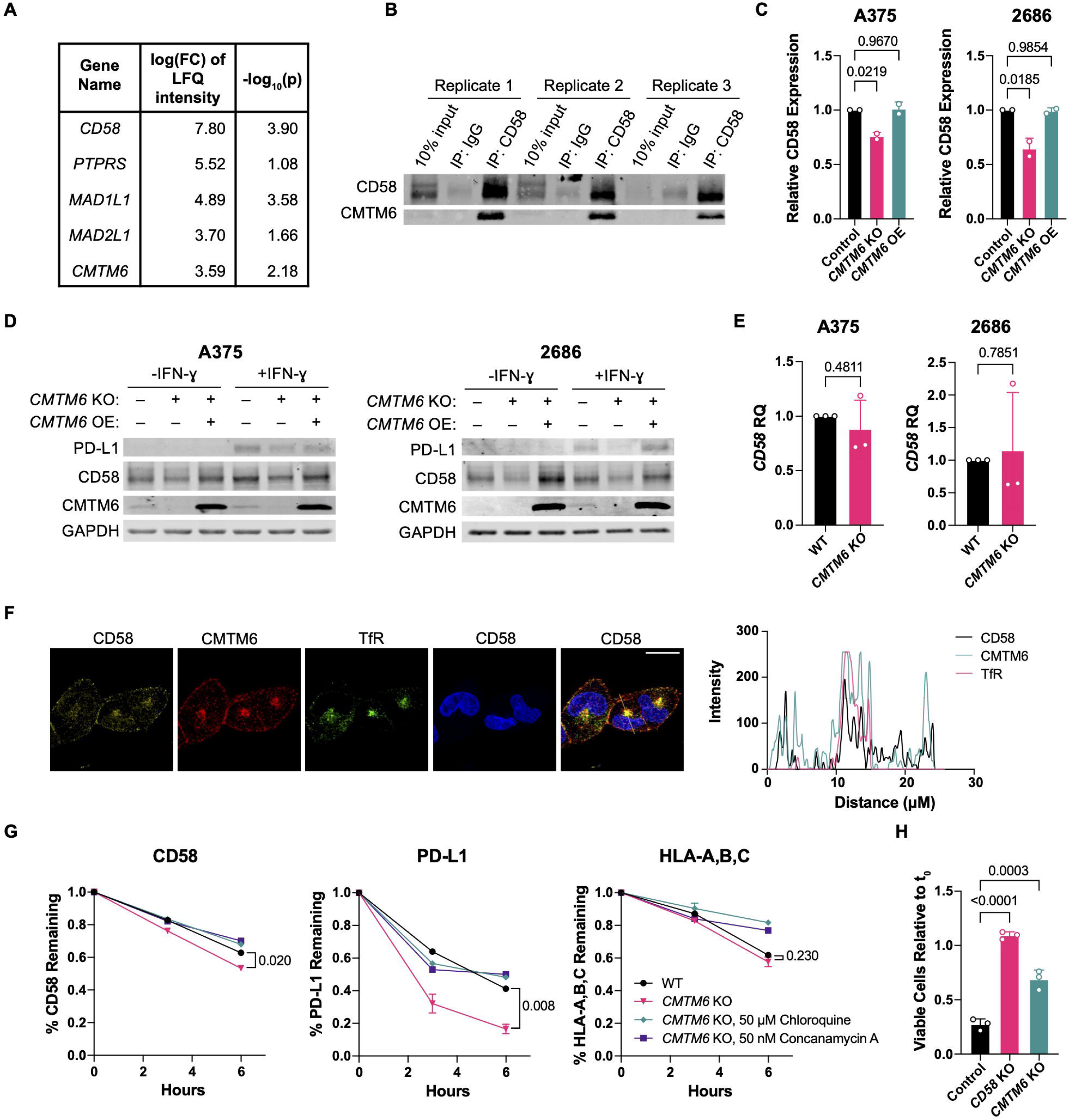
CMTM6 binds to and promotes protein stability of CD58 via endosomal recycling. (A) Genes whose encoded proteins were identified as interactors with CD58 by mass-spectrometry analysis of CD58 IP lysates from A375 melanoma cells. Inclusion criteria: FDR<0.05, log2 FC of LFQ intensity value >3, and an average spectral count within CRAPome database <2 to exclude background contaminants. Experiment was performed with three replicates. (B) Immunoblotting for CD58 and CMTM6 of A375 CD58 IP lysates used for MS-IP analysis. (C) Cell surface expression of CD58 in A375 and 2686 WT, *CMTM6* KO, and *CMTM6* OE cells as assessed by flow cytometry. (D) Immunoblotting for PD-L1, CD58, and CMTM6 in A375 and 2686 WT, *CMTM6* KO, and *CMTM6* (E) OE cells after 72 h with or without 10 ng/mL IFN-ɣ. (F) Relative gene expression of *CD58* in A375 and 2686 WT or *CMTM6* KO cells. (G) 2686 WT cells were fixed and immunostained for CD58, CMTM6, and transferrin receptor (TfR), a marker of recycling endosomes, and analyzed by confocal microscopy. Profile plots of relative fluorescence intensity along yellow line shown at right. Scale bar = 20 µm. 2686 WT and *CMTM6* KO cells were stained for cell surface CD58, PD-L1, and HLA-A,B,C with fluorophore-conjugated antibodies following 72 h with 10 ng/mL IFN-ɣ stimulation, and then incubated at 37 ºC for 3-6 h to allow for recycling of antibody-bound cell surface proteins in the presence or absence of lysosomal inhibitors chloroquine or concanamycin A, after which remaining CD58, PD-L1, and HLA-A,B,C expression was assessed by flow cytometry. Fold change in viable MaMel134 control, *CD58* KO, and *CMTM6* KO cells after 48 h co-culture with autologous TILs. (H) Representative blots shown from two independent experiments each (D). Representative images shown from two independent experiments (F). Experiments in (E) each performed with four technical replicates, with three independent experiments shown. Experiments in (C) performed in duplicate, with independent experiments shown. Experiments in (G) performed in duplicate, with representative experiment shown from two independent experiments. Statistical analysis performed using one-way ANOVA with Tukey’s multiple comparisons test (C,H) and two-sided T-tests (E,G). Data represent mean ± SD.

Notably, neither in the CRISPR/Cas9 nor in the MS-IP screen did we identify a direct genetic or protein interaction between CD58 and PD-L1, suggesting that their co-regulation is controlled by an alternative mediator. Because CMTM6 was nominated as a regulator of CD58 in both our CRISPR-Cas9 and MS-IP screens, and given its involvement in PD-L1 maintenance, we chose to investigate its role as such a potential regulator further. First, we validated that CMTM6 co-immunoprecipitates with CD58 in protein lysates used for MS-IP (Fig. 5B). Second, we generated *CMTM6* KO melanoma cell lines and showed that *CMTM6* loss led to decreased CD58 surface and total protein abundance in multiple models compared to otherwise isogenic controls (Fig. 5C-D, Fig. S6C). Rescue of *CMTM6* KO with a *CMTM6* ORF was sufficient to recover expression of protein CD58. In contrast, *CD58* mRNA levels were unaffected by perturbations of *CMTM6* with or without culturing with IFN-ɣ, demonstrating that CMTM6 regulates CD58 on a protein level but not a transcript level (Fig. 5E, Fig. S6D).

Given the role of CMTM6 in shuttling PD-L1 to the recycling endosome, we next investigated the subcellular localization of both CD58 and CMTM6. Immunofluorescence (IF) confocal imaging revealed that CD58 colocalizes with transferrin receptor (TfR), a marker of recycling endosomes, and additionally with CMTM6 in both recycling endosomes and the cell membrane (Fig. 5F, Fig. S6E). Furthermore, CMTM6 is known to shuttle PD-L1 to recycling endosomes to prevent its degradation in lysosomes, and so we reasoned that CMTM6 might regulate CD58 similarly. Indeed, in a flow-cytometry based degradation assay, we find that both CD58 and PD-L1, but not MHC Class I control, were degraded at higher rates in *CMTM6* KO compared to isogenic parental cells, and that this degradation could be restricted by different lysosome inhibitors (Fig. 5G). We therefore propose that, similar to its stabilization of PD-L1, CMTM6 binds to CD58 and promotes its recycling from the cell surface to endosomes, thus preventing its lysosomal degradation. In line with this finding, another positive regulator of CD58 that we had identified in our CRISPR/Cas9 KO screen was *DNAJC13*, whose encoded protein plays a role in mediating membrane trafficking from early to recycling endosomes^31^.

Functionally, *CMTM6* loss was sufficient to confer resistance to T cell-mediated tumor lysis in co-culture models, likely due to decreased CD58 levels, though not to the same extent as *CD58* loss (Fig. 5H). This limited resistance phenotype may be explained by either residual protein CD58 and/or loss of PD-L1 stability in *CMTM6* KO cells. In support of this, loss of CMTM6 resulted in reduced surface and total protein levels of PD-L1 (Fig. 5D, Fig. S6C,F), as previously reported^29,30^

### CMTM6 is a required mediator of concurrent PD-L1 expression in *CD58* null cells

Given the role of CMTM6 in regulating both PD-L1 and CD58, we reasoned that rather than interacting directly, PD-L1 and CD58 co-regulation might be mediated by CMTM6. To begin testing this hypothesis, we first performed co-IP of CMTM6 or CD58 pulldowns followed by probing for CMTM6, CD58 and PD-L1. While CMTM6 binds to both CD58 and PD-L1 (Fig. 6A), CD58 binds to CMTM6, but not PD-L1 (Fig. 6B). Furthermore, *CD58* loss had no impact on *CD274* (PD-L1) mRNA expression, indicating that CD58 regulates PD-L1 only at a protein level (Fig. 6C, Fig S7A). Interestingly, loss of *CD58* resulted in increased co-immunoprecipitation of CMTM6 and PD-L1, while *CD58* OE resulted in reduced CMTM6/PD-L1 binding (Fig. 6D). We hypothesized that CD58 regulates PD-L1 expression by modulating CMTM6/PD-L1 binding, and that with higher expression of *CD58*, additional CD58 may compete for CMTM6 and therefore result in increased PD-L1 degradation. To test this, we rescued CD58 KO cells with *CD58-TM* OE and measured the rate of degradation of PD-L1 (**Methods**). We find that forced *CD58* expression resulted in a more rapid decline of PD-L1 protein levels, which was rescued by addition of lysosome inhibitors (Fig. 6E). Together, these results suggest that loss of *CD58* results in increased PD-L1 levels by stabilizing CMTM6/PD-L1 interactions and therefore reducing PD-L1 degradation in the lysosome.

**Figure 6:**
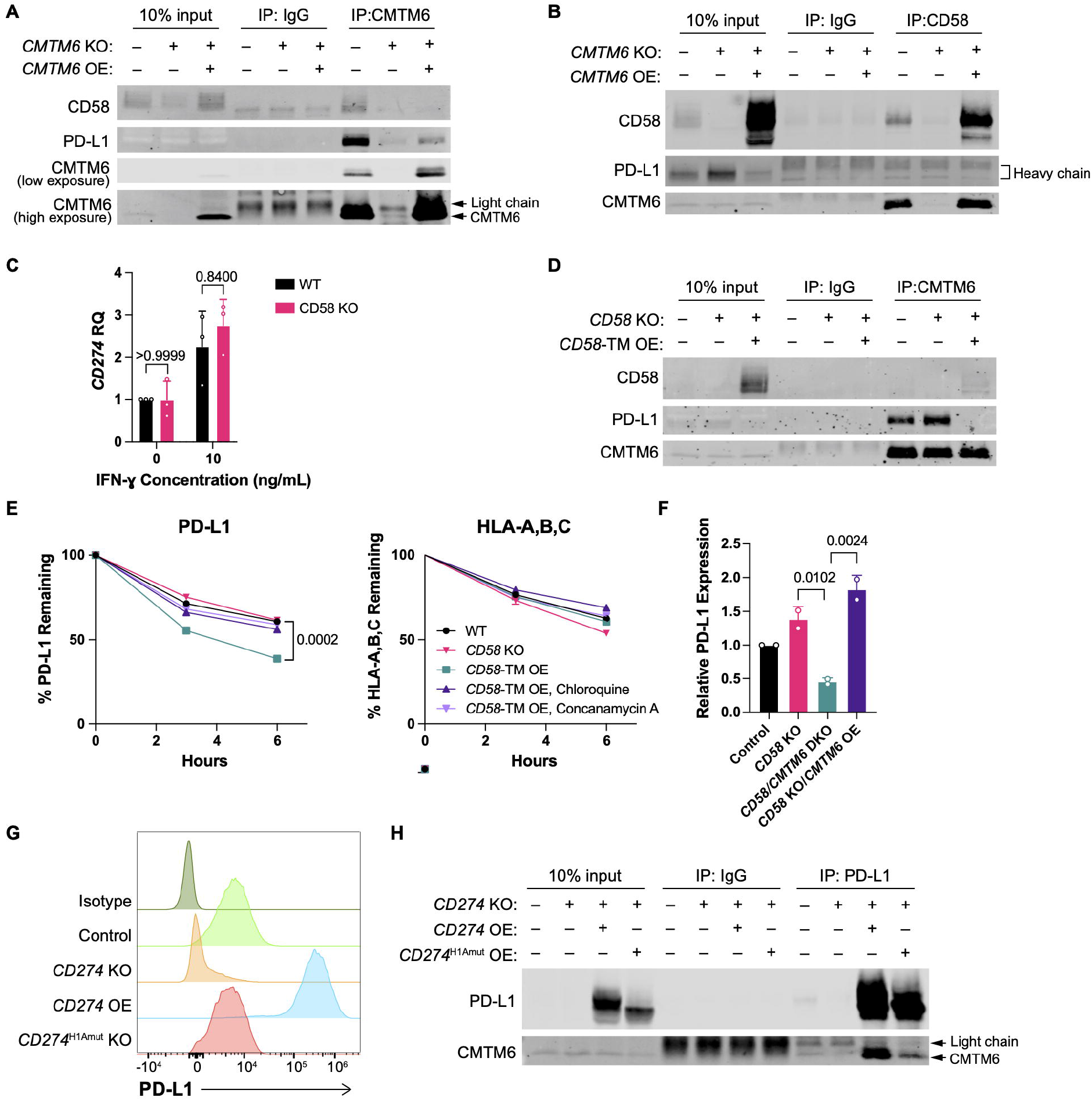
CMTM6 is a necessary mediator of CD58’s regulation of PD-L1. (A) Co-immunoprecipitation of PD-L1 and CD58 with CMTM6 pulldown in 2686 WT, *CMTM6* KO, and *CMTM6* OE cells after 72 h with 10 ng/mL IFN-ɣ. (B) Co-immunoprecipitation of CMTM6, but not PD-L1, with CD58 pulldown in 2686 WT, *CD58* KO, and *CD58* OE cells after 72 h with 10 ng/ml IFN-ɣ. (C) Relative gene expression of *CD274* in 2686 WT and *CD58* KO cells after 72 h with or without 10 ng/ml IFN-ɣ. (D) Co-immunoprecipitation of CD58 and PD-L1 with CMTM6 pulldown in 2686 WT, *CD58* KO, and *CD58* OE cells after 72 h with 10 ng/mL IFN-ɣ. (E) Remaining PD-L1 and HLA-A,B,C in 2686 WT, *CD58* KO, and *CD58*-TM OE cells (pre-stimulated with 10 ng/mL IFN-ɣ for 72 h) after 3 and 6 hours of incubation at 37 ºC post-staining. (F) Surface expression levels of PD-L1 in 2686 control, *CD58* KO, *CD58/CMTM6* DKO, and *CD58* KO/*CMTM6* OE cells after 72 h with 10 ng/mL IFN-ɣ as assessed by flow cytometry. (G) Surface expression of PD-L1 in 2686 control, *CD274* KO, *CD274* OE, and *CD274*^*H1Amut*^ OE cells after 72 h 10 ng/mL IFN-ɣ. Counts normalized to mode. (H) Co-immunoprecipitation of CMTM6 with PD-L1 pulldown in 2686 control, *CD274* KO, *CD274* OE, and *CD274*^*H1Amut*^ OE cells after 72 h 10 ng/mL IFN-ɣ. Representative blot shown from two independent experiments (A,B,D,H). Experiments in (C) performed with four technical replicates, with independent experiments shown. Experiments in (E) performed in duplicate, with representative experiment shown from two independent experiments. Experiments in (F) performed in duplicate, with two independent experiments shown. Statistical analysis performed using two-sided T-test (C, E) and one-way ANOVA with Tukey’s multiple comparisons testing (F). Data represent mean ± SD.

To determine if CMTM6 is in fact necessary for upregulation of PD-L1 with *CD58* loss, we generated *CD58*/*CMTM6* double knockout (DKO) cell lines (Fig. S7B). While *CD58* KO resulted in increased PD-L1 expression, *CD58*/*CMTM6* DKO did not; in contrast, rescue of *CD58*/*CMTM6* DKO with *CMTM6* OE reconstituted concurrent PD-L1 upregulation (Fig. 6F, Fig. S7C), demonstrating that *CMTM6* is necessary for increased PD-L1 levels in *CD58* loss. Lastly, we sought to determine the CMTM6 binding site in PD-L1. A prior study demonstrated that an N-terminal domain (spanning amino acids (aa) 20-32) was necessary for stabilization of PD-L1 at the cell membrane, and that addition of an antibody targeting this epitope (clone H1A) reduced PD-L1 levels^32^. This domain therefore represents a putative binding site for CMTM6. To test this, we generated PD-L1/*CD274* KO melanoma cells and rescued them with either WT *CD274* or a mutated form of *CD274*, in which the coding region for aa 20-32 was scrambled (PD-L1/*CD274*^*H1Amut*^). Overexpression of PD-L1/*CD274*^*H1Amut*^ failed to rescue surface expression of PD-L1 (Fig. 6G, S7D), while total protein levels of PD-L1 were ∼4 fold higher in PD-L1/*CD274*^*H1Amut*^ cells compared to parental cells (S7E-F). Furthermore, PD-L1/*CD274*^*H1Amut*^ demonstrated significantly impaired binding to CMTM6 compared to PD-L1 in WT PD-L1/*CD274* OE cells (Fig. 6H), suggesting that reduced surface PD-L1 in PD-L1/*CD274*^*H1Amut*^ cells was due to impaired CMTM6 binding to PD-L1, and therefore increased degradation.

Together, these data support a model in which both CD58 and PD-L1 require CMTM6. When *CD58* expression is lost, additional PD-L1 protein binds to released CMTM6 and is stabilized at the cell surface through reduced lysosomal degradation (Fig. S7G).

## DISCUSSION

Here, we find that an intact CD58:CD2 axis is necessary for effective anti-tumor immunity, and that its absence results in immune resistance via multiple mechanisms, including impaired T cell activation, inhibited entry and proliferation within tumors, and concurrent upregulation of PD-L1. We demonstrate that these mechanisms are orthogonal to clinically validated mechanisms of resistance to immunotherapies, such as impaired antigen presentation^4,5^ or IFN-ɣ-JAK/STAT signaling^3,33^. In addition to melanoma^7^, we also show that these mechanisms may play a role in kidney cancer patients treated with immune checkpoint blockade^21^. Furthermore, *CD58* loss has previously been implicated in immune escape by hematological cancers^34-37^ and has recently been identified as a mechanism of resistance to CD19-targeting CAR-T cells in patients with diffuse large B cell lymphoma (DLBCL)^20^. Together, this evidence suggests that cancer cell intrinsic expression of *CD58* may play an important role in cancer immune evasion across different lineages and types of immunotherapies; thus, our study provides important mechanistic insights resolving the underpinnings of this immune axis.

Expression of the primary co-stimulatory protein CD28 on T cells is required for response to anti-PD1 therapy^16^; however, we find that TILs in most patients lose *CD28* expression across differentiation states, thus contributing to anti-PD-1 resistance. In this context, CD58 is the most effective costimulatory ligand in CD8+CD28-T cells^19^. We show that virtually all TILs in patients with both melanoma and kidney cancer maintain *CD2* expression, and that *ex vivo* expanded human CD28-CD8+ TILs can achieve polyfunctionality and potent anti-tumor activity upon stimulation with recombinant CD58 protein. Thus, the CD58:CD2 axis may represent a “last resort” for T cell activation. Furthermore, restoration of this axis may serve as a promising therapeutic approach in tumors that acquire immune resistance through cancer cell intrinsic *CD58* downregulation, as CD2 co-stimulation may mobilize a major pool of TILs that is largely untapped by existing cancer immunotherapies. Such therapeutic strategies require careful design to prevent non-specific effects. A recent study has found that engineered CAR-T cells coupled with a CD2 receptor may overcome resistance due to *CD58* loss in an *in vivo* model of DLBCL^20^. In solid tumors, trispecific antibodies were recently shown to be effective in targeting CD28+ T cells against HER2+ breast cancer *in vivo*^38^; similar antibodies might be engineered to instead stimulate T cells via CD2 in a targeted manner. The utility of CD2 co-stimulation could be expanded to CD28+CD8+ T cell populations as well, as recent work demonstrates that CD2 and CD28 co-stimulation are comparable in their ability to promote proliferation, cytokine production, and effector differentiation in naïve T cells, though CD2 stimulation more efficiently maintains a pool of progenitor-like T cells that are important for response to PD-1 blockade^39^.

How cancer cells balance potential co-stimulatory signals (which in certain contexts may confer a fitness advantage^40^) and co-inhibitory signals remains obscure, but is fundamental to our understanding of tumor-immune interactions and the efficacy of cancer immunotherapies. Here, we show that loss of *CD58* results in increased PD-L1 protein abundance, thus leading to a dual setback to T cell mediated anti-tumor immunity: lack of stimulation by CD58 and increased inhibition by PD-L1. Through unbiased CRISPR-Cas9 KO and MS/IP screens, we identify CMTM6, which was previously shown to be required for PD-L1 maintenance^29,30^, as a key regulator of CD58 protein. We show that CD58 and PD-L1 are co-regulated by CMTM6, rather than interacting directly, and we demonstrate that CMTM6 is necessary for concurrent upregulation of PD-L1 in the context of *CD58* loss. Our data support a model wherein CD58 and PD-L1 compete for CMTM6 for protein maintenance in the cell membrane and recycling endosomes, thus preventing their lysosomal degradation. Interestingly, PD-1:PD-L1 engagement has itself been shown to be localized at the corolla formed by CD58:CD2 ligation in the immune synapse in model systems, thereby subsequently decreasing CD2 signal amplification in T cells^41^. Thus, CD58’s regulation of PD-L1 further helps to maintain intact CD58:CD2 axis signaling, as it prevents buffering of this interaction by PD-1:PD-L1 ligation. Importantly, we also show that a short N-terminal domain of PD-L1 is necessary for CMTM6 binding, and that disruption of this domain results in destabilization and degradation of PD-L1. Antibodies targeting this domain – for example, H1A^32^ – therefore represent another therapeutic opportunity in the context of ICB. This is particularly tractable, as our human data suggest that tumors with resistance to ICB exhibit downregulation rather than genetic loss of *CD58*; *CD58* expression is therefore potentially rescuable as a therapeutic substrate.

In patients with melanoma and non-small cell lung cancer, CMTM6 protein abundance was associated with increased T cell abundance in pre-treatment biopsies^42,43^, but its role as a predictive biomarker remains unclear^44^, likely due to its dual role in co-regulating PD-L1 and CD58. Recent studies indicate a possible role of CMTM6 in genome stability and the mTOR pathway, and suggest that inhibition (or degradation) of total CMTM6 may reduce tumor severity and metastatic behavior^45,46^. Our study highlights a complex role of CMTM6 in both pro- and anti-tumor immunity regulation; thus, focusing on specific interactions may be important in the development of therapeutics targeting CMTM6.

To our knowledge, this is one of the first mechanistic descriptions of how co-inhibitory and costimulatory proteins are co-regulated in cancer. While we focused on the interaction of CD58 and PD-L1, this raises the broader conceptual question of how perturbation (either pharmacological or genetic) of one checkpoint influences that of others in the same cell. Systematic approaches, such as combinatorial Perturb-CITE-seq of human models^8^ may provide important insights and inform rationale for the next generation of combinations of cancer immunotherapies.

In summary, our work contributes three major advances. First, we identify a central and necessary function of the CD58:CD2 axis in anti-tumor immunity and demonstrate that defects in this interaction confer resistance to T cell-mediated killing by multiple mechanisms. Second, we identify CMTM6 as a critical regulator in the yin and yang of co-stimulatory and co-inhibitory cues by cancer cells. Third, we present the basis for potential therapeutic development to overcome resistance caused by *CD58* downregulation or loss, which may be relevant across diseases and immunotherapies.

## Acknowledgments

P.H., Z.W., L.C., M.M., and E.K.M. are supported by Training Grant T32GM007367. B.I. is supported by National Institute of Health/National Cancer Institute (NIH/NCI) grants K08CA222663, R21CA263381, and R37CA258829; the American Cancer Society Research Scholar Grant 21-104-01-IBCD; the Burroughs Wellcome Fund Career Award for Medical Scientists; a Tara Miller Melanoma Research Alliance Young Investigator Award; the Louis V. Gerstner, Jr. Scholars Program; and the V Foundation Scholars Award. K.W.W. is supported by NIH grants R01CA238039 and R01CA25159. C.A. was supported by a TL1 award. Results shown here are in part based upon data generated by the TCGA Research Network: https://www.cancer.gov/tcga. This work was supported by Herbert Irving Comprehensive Cancer Center core facility grants (P30CA013696, S10OD020056).

## Author Contributions

B.I. supervised the work. P.H. designed, performed, or participated in performing all experiments and analyses. J.C.M. and M.R. designed and performed experiments. C.J.F. performed analyses of CRISPR-Cas9 screens. S.B.S., Z.W., O.K., A.D.A., L.C., B.T.F., R.S., C.R.A., M.M., H.F., and E.K.M. performed experiments. J.B., Y.W. and R.S. performed analyses of single-cell data and MS screens. N.V., S.F.B., S.L.R., C.B., E.M.M., K.W.W., D.S. and G.K.S. provided critical reagents, materials, and resources. P.H. and B.I. wrote the manuscript.

## Declaration of Interests

B.I. is a consultant for Volastra Therapeutics Inc., Johnson & Johnson/Janssen, and received honoraria from AstraZeneca and Merck. None of these represent a conflict of interest pertaining to the presented work. K.W.W. serves on the scientific advisory board of T-Scan Therapeutics, SQZ Biotech, Nextechinvest and receives sponsored research funding from Novartis. He is a co-founder of Immunitas, a biotech company. These activities are not related to the research reported in this publication. P.H., J.C.M. and B.I. filed a patent pertaining to the presented work.

**Figure S1**: **RNA-seq analysis of melanoma patient tumors. Related to Fig. 1**.

(A) Comparison of *CD58* expression (shown by normalized count) in cutaneous and uveal melanomas from TCGA RNASeqV2 data, computed by RSEM.

(B) *CD58* expression in cancer cells from scRNA-seq data from previously published dataset of eight renal cell carcinoma tumors; five patients were exposed to ICB targeting the PD-1 axis and three patients were treatment naïve^21^.

(C) scRNA-seq expression data from a previously published dataset of 33 human melanoma tumors^6^, split into *CD8*+*TCF7*+ and *CD8*+*TOX*+ T cells. Samples are also separated by treatment status into either treatment-naïve tumors or a group of 15 post-ICB resistant tumors.

(D) As in (B), but analysis is performed on another previously published scRNA-seq dataset of 46 tumor biopsies of metastatic melanoma patients treated with anti-PD-1 therapy^22^. Samples are separated into *CD8*+*TCF7*+ and *CD8*+*TOX*+ T cells, and also by patient clinical response to therapy.

(E) *CD28* and *CD2* expression among CD8+ T cells from same RCC tumor dataset as in (B).

**Figure S2**: **CD58-CD2 ligation is critical to T-cell-mediated killing across cell lines. Related to Fig. 1**.

(A) Surface CD58 expression in 2686 DsRed, MaMel134 DsRed, and A375 control, *CD58* KO, and *CD58*-TM or *CD58*-GPI OE cells used in co-culture and flow cytometry experiments.

(B) Widefield images of 2686 and MaMel134 melanoma/TIL co-cultures shown in Figure 1A. Arrowheads indicate T cell clusters following T cell activation. Scale bar = 100 µm.

(C) PBMC-derived T cells were sorted for CD8+ T cells, which were then engineered to express a NY-ESO-1-specific TCR carrying an HA tag. CD8 and HA expression on engineered T cells was assessed by flow cytometry.

(D) Fold change in number of viable cells after 24 h co-culture of 2686 patient melanoma control, *B2M* KO, and *CD58* KO cells with NY-ESO-1-specific TCR engineered T cells.

(E-F) Fold change in number of viable MaMel134 cells after 48 h co-culture with autologous TILs in the presence of anti-CD2 blocking antibody (LoCD2b) (E), anti-CD58 (TS2/9) blocking antibody (F), or IgG isotype control.

(G-H) Binding of anti-CD58 antibody clones TS2/9 (G) and 1C3 (H) to cell surface CD58 in MaMel134 and 2686 control, *CD58* KO, *CD58-*TM, *CD58*^*K34A*^-TM, *CD58*-GPI, or *CD58*^*K34A*^-GPI OE cells as assessed by flow cytometry. Clones TS2/9 and 1C3 are known to inhibit CD58:CD2 binding. Counts normalized to mode.

(I) IFN-ɣ concentrations from cleared co-culture media from experiments shown in Figure 1C.

(J) Surface CD2 expression of WT and *CD2* KO TILs assessed by flow. Counts normalized to mode.

(K-L) MaMel134 TILs were stained with CFSE and stimulated for four days with 1 µg/mL anti-CD3 (OKT3) antibody ± 1 µg/mL anti-CD28 or CD58-Fc chimera. Number of cell divisions was quantified by flow cytometry analysis of remaining CFSE staining (K) and IL-2 concentrations within cleared culturing media were quantified by ELISA (L).

(M) Surface CD48 expression in A375, MaMel134, 2686, and control Jurkat WT cells. Counts normalized to mode.

Representative images shown in (B) from at least two independent experiments each. Experiments performed in duplicate (K-L) or triplicate (D-F,I), with representative experiment shown from at least two independent experiments each. Statistical analysis performed using one-way ANOVA with Tukey’s multiple comparisons testing (D-F,I,L). Data represent mean ± SD.

**Figure S3: *In vivo* model of ACT-treated melanoma recruits intratumoral CD8+ TILs. Related to Fig. 2**.

(A) Example IL-2 serum concentration in NOG mice (n=8).

(B) Flow gating scheme used for analyzing processed mouse tumors. Fluorescence minus one (FMO) controls were used to determine Ki-67 and CD28 expression.

(C) Percent of CD8+ cells among MaMel134 TILs used for each round of ACT.

(D-E) Percent of CD8+ cells among hCD45+ cells within control v. *CD58* KO melanoma tumors (D) and control v. *CD58* OE tumors at experiment endpoint (E).

(F) CD8:mCD45+ ratio within ACT-treated control and *CD58* KO tumors plotted against final tumor weight. Best fit semilog line shown. R^2^ 0.1832, 0.3376, respectively.

(G-H) Percent of CD8+ TILs within ACT-treated control and *CD58* OE (G) or *CD58* KO (H) cells that do not express CD28, as analyzed by flow cytometry.

Statistical analysis performed using paired two-sided T-tests (D-E, G-H). Line at median.

**Figure S4: CD58 regulates PD-L1 expression across multiple cell lines. Related to Fig. 3**.

(A) Surface PD-L1 expression in A375 WT, *CD58* KO, *CD58*-TM OE, and *CD58-*GPI OE cells after 72 h with 10 ng/mL IFN-ɣ.

(B) Whole protein CD58 and PD-L1 expression in A375 WT, *CD58* KO, *CD58-TM*, and *CD58-GPI* OE cells after 72 h in the absence or presence of 10 ng/mL IFN-ɣ.

(C) Surface HLA-A,B,C expression in A375 WT, *CD58* KO, *CD58-TM*, and *CD58-GPI* OE cells.

(D) Surface PD-1 expression among CD8+ MaMel134 TILs after overnight stimulation with or without 1 µg/mL anti-CD3 (OKT3) prior to co-culture with autologous melanoma cells shown in Figure 3E. Counts normalized to mode.

Experiments (A,C) performed in duplicate two or three independent experiments shown. Representative blot shown in (B) of at least two independent experiments. Statistical analyses performed using one-way ANOVA with Tukey’s multiple comparisons test. Data represent mean ± SD.

**Figure S5: Preparation and analysis of CRISPR/Cas9 KO screen to identify regulators of CD58. Related to Fig. 4**.

(A) Cas9 activity of A375 cells transduced with pLX_311-Cas9 lentivirus was assessed by transduction with pXPR-011 lentivirus encoding both eGFP and eGFP-targeting sgRNA. Percent Cas9 activity represented by percentage of eGFP-cells.

(B) A375 Cas9-expressing cells were transduced with Brunello genome-wide sgRNA library at increasing virus volumes per 1 million cells for a desired transduction efficiency of 20% (MOI<0.3). 30 µL virus was used for screen.

(C) Anti-CD58-APC antibody was titrated for use in assessing CD58 surface expression in CRISPR/Cas9 screen. 2 µg/mL antibody was used for FACS.

(D) Following 10 days of culturing, CRISPR/Cas9 screen cells were sorted based on CD58 surface expression. Representative gating of CD58^lo^ (P6) and CD58^mi^ (P5) control populations shown.

(E) Following sort of target and control populations, sgRNA sequences were PCR amplified, with a desired PCR product size of ∼354 bp (arrow).

(F) Distribution of sgRNA guide enrichment for top 20 gene targets (ranked by P-value) for each replicate of CRISPR/Cas9 screen (Table S1-2). Red bars indicate log2 fold enrichment of each sgRNA per gene within the CD58^lo^ population compared to CD58^mi^ control population for each replicate (biological replicates 1 and 2, each with technical replicates A and B), with genes ranked from top to bottom by P-value.

**Figure S6: CMTM6 regulates CD58 at a protein level across multiple cell lines. Related to Fig. 5**.

(A) Genes whose encoded proteins were identified as interactors with CD58 by mass-spectrometry analysis of CD58 immunoprecipitates from 2686 melanoma cells. Experiment performed with three biological replicates.

(B) Immunoblotting for CD58 of 2686 CD58 immunoprecipitates used for MS-IP analysis.

(C) Whole protein PD-L1, CD58, and CMTM6 expression in MaMel134 DsRed and WM852 control, *CMTM6* KO, and *CMTM6* OE cells after 72 h ± 10 ng/mL IFN-ɣ stimulation.

(D) Relative gene expression of *CD58* in A375 and 2686 WT or *CMTM6* KO cells after 72 h 10 ng/mL IFN-ɣ stimulation.

(E) 2686 WT cells were fixed and immunostained for CD58 alongside markers of ER (Calnexin), Golgi (GM130), early endosomes (EEA1), lysosomes (LAMP1), and recycling endosomes (TfR) and analyzed by confocal microscopy. Profile plots of relative fluorescence intensity along yellow line shown at right. Scale bars = 20 µm.

(F) Flow cytometry analysis of surface PD-L1 in A375 and 2686 WT, *CMTM6* KO, and *CMTM6* OE cells after 72 h 10 ng/mL IFN-ɣ stimulation.

Representative blots shown from at least two independent experiments each (C). Experiments in (D) each performed with four technical replicates, with three independent experiments shown.

Representative images shown from two independent experiments (E). Experiments in (F) performed in duplicate, with two independent experiments shown. Statistical analysis performed with two-sided T-test (D) and one-way ANOVA with Tukey’s multiple comparisons (F). Data represent mean ± SD.

**Figure S7: CMTM6 is critical for CD58’s regulation of PD-L1. Related to Fig. 6**

(A) Relative gene expression of *CD274* in A375 WT or *CD58* KO cells after 72 h 10 ng/mL IFN-ɣ stimulation.

(B) Validation of whole protein CD58 and CMTM6 expression levels by western blot of cells used in Figure 6F and S7C.

(C) Surface expression levels of PD-L1 in A375 WT, *CD58* KO, *CD58/CMTM6* DKO, and *CD58* KO/*CMTM6* OE cells after 72 h with 10 ng/mL IFN-ɣ.

(D-E) Relative surface (D) and whole protein (E) PD-L1 expression in 2686 control, *CD274* KO, *CD274* OE, and *CD274*^*H1Amut*^ OE cells after 72 h ± 10 ng/mL IFN-ɣ assessed by surface ± intracellular flow staining of PD-L1.

(F) Immunoblotting for PD-L1 expression in 2686 control, *CD274* KO, *CD274* OE, and *CD274*^*H1Amut*^ OE cells after 72 h ± 10 ng/mL IFN-ɣ.

(G) Proposed model of PD-L1 regulation by CD58. In WT conditions, CD58 and PD-L1 both bind to CMTM6, which stabilizes both proteins at the cell surface and prevents their degradation by lysosomes. In the absence of CD58, excess CMTM6 is available to bind to PD-L1 and increase its expression at the cell surface. Created with BioRender.

Experiments performed in (A) with four technical replicates, with three independent experiments shown. Experiments in (C-E) performed in duplicate, with two independent experiments shown. Representative blot shown (F) from two independent experiments shown. Statistical analysis performed using two-sided T-tests (A) and one-way ANOVA with Tukey’s multiple comparison testing (C-E). Data represent mean ± SD.

## METHODS

### Cell culture

A375 melanoma cells and HEK293T cells were purchased from ATCC, and WM852 melanoma cells were purchased from Rockland Inc. Patient-derived melanoma line 2686 and matched TILs were previously derived (under institutional review board protocol no. 2004-0069) and provided by MD Anderson Cancer Center. Patient-derived melanoma cell line MaMel134 and matched TILs were provided by UK-Essen. Jurkats were a gift from Adam Mor. A375 and HEK293T cells were cultured in DMEM medium supplemented with 10% heat-inactivated fetal bovine serum (D10). All other melanoma cell lines were cultured in RPMI 1640 medium supplemented with 10% heat-inactivated fetal bovine serum, GlutaMax, 10 mM HEPES, 10 mg/L insulin, 5.5 mg/L transferrin, 6.7 µg/L sodium, and 55 µM 2-mercaptoethanol (all from Thermo Fisher Scientific) at 37 ºC and 5% CO2 in a humidified incubator. TILs were cultured in T cell media (RPMI 1640 medium supplemented with 10% human AB serum (Fisher), GlutaMax, 10 mM HEPES, 100 IU/mL penicillin and 100 µg/mL streptomycin (Thermo)), supplemented with clinical grade 300-3000 IU/mL human IL-2 (Proleukin, Chiron). All cell lines were tested routinely for mycoplasma contamination using PlasmoTest (InvivoGen).

### Generation of NY-ESO-1 TCR T cells

Primary human cells were isolated from blood collars provided by Brigham and Women’s Hospital. Human blood mononuclear cells were isolated by Ficoll-Paque gradient centrifugation using SepMate isolation tubes (StemCell #85450). CD8^+^ T cells were then isolated by negative selection using the EasySep CD8 isolation kit (StemCell #17953) according to manufacturer’s instructions. T cells were activated with CD3/CD28 Dynabeads (Thermofisher #11132D) at a 1:1 ratio for 48 hours in media supplemented with 30U/mL of IL-2. NY-ESO-1 TCR was introduced as described previously (Roth et al., 2018; Matthewson et al., 2021). Briefly, the NY-ESO-1 construct was introduced into the endogenous *TRAC* locus by homology-directed repair. T cells were electroporated with a ribonucleoprotein complex (RNP) of Cas9 protein with a bound TRAC gRNA as well the NY-ESO-1 TCR construct. The TRAC gRNA (60mM) was assembled by incubating equimolar quantities of TRAC crRNA and tracrRNA at 95°C for 5 minutes. Then, the resulting gRNA was assembled into RNPs by incubation with an equal volume of Cas9-NLS protein (20mM) at 37°C for 15 min. CD8 T cells (1×10^6^) were electroporated with Cas9-TRAC gRNA RNPs and 2.5mg of single-stranded NY-ESO-1 TCR DNA template. The ssDNA template was synthesized using GuideIt Long ssDNA kit (Takara #632666) according to the manufacturer’s instructions. Electroporated T cells were cultured overnight and then re-activated with Dynabeads at a 1:1 ratio. T cells were expanded for 10 days in media containing IL-2 and then sorted based on expression of CD3 and NY-ESO-1 TCR using the HA epitope tag attached to the N-terminus of the mature NY-ESO-1 TCR a chain. Sorted T cells were expanded, frozen in aliquots, and stored at -80°C until use.

### Expansion of autologous TILs or NY-ESO-1 TCR T cells

Patient-derived TILs and human CRISPR-Cas9 NY-ESO1 TCR knock-in CD8+ T cells were expanded using a rapid-expansion protocol (REP) as previously described^8^. For expansion, PBMCs were isolated from three different donors and used as feeder cells as follows. After isolation of PBMCs via Ficoll gradient centrifugation (Ficoll-Paque Premium, Cytiva), red blood cells were lysed using ACK-lysis buffer (Thermo), and PBMCs were washed once in PBS and then resuspended in AIM-V media (Thermo) before irradiating with 5,000 rad. On day 0, 5e^5^ – 1e^6^ TILs or edited CD8 T cells were then added to 100e^6^ irradiated feeder cells in a G-Rex 10 bottle (Wilson Wolf) in REP media (1:1 mixture of AIM-V media and T cell media) supplemented with a final concentration of 30 ng/mL anti-CD3 antibody (OKT3) (Miltenyi) and 3,000 IU/mL human recombinant IL-2 (Proleukin, Chiron). On day 2, 3,000 IU/ml human recombinant IL-2 was added. On day 5, fresh media and IL-2 were added by half-media exchange. On day 7, cells were counted and >60e^6^ cells were propagated to a G-Rex 100 bottle in a total volume of 600 mL AIM-V with 3,000 IU/mL IL-2. On day 10, additional 400 mL AIM-V with 3000 IU/mL IL2 was added. On day 12 additional 3000 IU/mL IL-2 were added. On day 14 expansion was complete and the expanded cells were collected, cryopreserved, and stored in liquid nitrogen until use in *in vitro* or *in vivo* experiments.

### CRISPR-Cas9 KO cell line generation

Virus-free KO human melanoma cell lines and TILs were generated by nucleofection of Cas9 ribonucleoproteins (RNPs). TILs were pre-stimulated on culture dishes coated with 100 ng/mL anti-CD3 (OKT3) antibodies overnight. Target sequences were selected from CRISPick^27,47^; the following crisprRNA (crRNA, IDT) sequences were used: AATGCTCTGGTATCATGCAT (*CD58*), ACTGCTTGTCCAGATGACTT (*CD274*), GGTGTACAGCCCCACTACGG (*CMTM6*), CGATGATCAGGATATCTACA (*CD2*). Briefly, equimolar ratios of crRNA and trans-activating crispr RNA (tracrRNA) were incubated at 95 ºC for 5 min and then cooled to room temperature to form guideRNA (gRNA). Recombinant Cas9 enzyme (MacroLab) was incubated with gRNA at a 1:10 molar ratio at 37 ºC for 15 min to form RNP complexes. Melanoma cells were resuspended at a cell density of 50,000 cells in 20 µL SF electroporation buffer with supplement (Lonza) and TILs were resuspended at a cell density of 1 million cells in 20 µL P3 electroporation buffer with supplement (Lonza) and then combined with 3 µL of RNP mixture and nucleofected using program DJ-110 (melanoma) or EH100 (TILs) on a 4D Nucleofector (Lonza). Melanoma cells were immediately recovered in full melanoma media in 12-well plates. TILs were left to incubate for 15 min at 37 ºC prior to transfer to a 24 well recovery plate containing 300 IU/mL hIL-2. Pure cancer KO populations were generated by cell surface staining and sorting by FACS as described below. CD2 editing in TILs achieved >95% editing and required no sorting. Prior to coculture, TILs were rested for 10 days and unedited TILs from the initial stimulation step were cultured in parallel and used as controls.

### *CD58, CMTM6*, and *PD-L1* OE construct cloning

*CD58, CMTM6, PD-L1, and PD-L1*^*H1Amut*^ open reading frames (ORFs) with a 5’ Kozak sequence were flanked by *att*B cloning sites and ordered as gene blocks (GeneWiz), which were then cloned into pDONR™221 (Thermo) to create an entry clone using the Gateway Cloning system (Thermo). Donor constructs were then cloned into the constitutive expression vector pLEX_307 (gift from David Root; Addgene #41392) or pLX_TRC311 (gift from John Doench; Addgene #113668). *CD58*^*K34A*^ OE plasmids were generated through site-directed mutagenesis of existing *CD58* ORF plasmids using the QuikChange XL Site-Directed Mutagenesis Kit (Agilent Technologies) per manufacturer protocol. Forward primer: TTTTCCAGTTCTGCAACTGCATCCTTTTGTTTTTTCCATAGGACCTCTTTTAA. Reverse primer: TTAAAAGAGGTCCTATGGAAAAAACAAAAGGATGCAGTTGCAGAACTGGAAAA. Plasmid insert validation was performed via Sanger sequencing (GeneWiz). Whole plasmid sequencing validation of empty vectors was performed by the Massachusetts General Hospital Center for Computation and Integrative Biology DNA Core. All plasmid design and Sanger sequencing analysis was performed using Snapgene 6.0 (GSL Biotech LLC, San Diego).

### Lentivirus production

HEK293T cells were seeded in 6-well plates in D10 media. The following day, 3 µL TransIT-LT1 transfection reagent (Mirus Bio) was mixed with 15 µL Opti-MEM (Thermo) and incubated for 5 min at room temperature (RT). A mixture of 500 ng expression plasmid, 500 ng packaging plasmid psPAX2 (gift from Didier Trono, Addgene #12260), and 250 ng envelope plasmid pMD2.G (gift from Didier Trono, Addgene #12259) was prepared to a final volume of 37.5 µL in Opti-MEM. Transfection reagent mix was combined with plasmid mix, incubated for 30 min at RT, and then added dropwise to HEK293T cells. After 24 h, cell media was replaced with DMEM supplemented with 20% FBS. Cell media was collected and replaced after 48 h and 72 h and filtered through a 0.45 µm syringe filter (Thermo). Lentivirus was stored at -80 ºC.

### Transduction of *CD58, CMTM6*, and *PD-L1* ORF constructs

Human melanoma cells were seeded in 6-well plates with 0.1-1 mL lentivirus in a final volume of 3 mL with 4 µg/mL polybrene (Millipore). Cells were spun at 1,000 x g for 2 h at 30 ºC, and an additional 3 mL media was added to each well after spinning. Cells were then incubated overnight at 37 ºC and 5% CO2 in a humidified incubator. The next day, infected and non-infected cells were seeded across two wells each in a 6-well plate at equal seeding density to monitor transduction efficiency, with all remaining cells grown up in appropriate sized flasks. 2 days after transduction, one well each of cells was put into puromycin or blasticidin selection (Thermo). Once all non-infected cells were dead, transduction efficiency was calculated as the ratio of the number of infected cells surviving in selection divided by the number of infected cells growing without selection.

### Co-culture assays

Melanoma target cells expressing nuclear localization signal (NLS)-dsRed were co-cultured with matched TILs or NY-ESO-1-targeting engineered T cells as previously described^8^. Briefly, melanoma target cells were seeded in a black-walled 96-well plate (Corning) at a cell density of 10,000 cells/well. After 16 h, culture medium was replaced with 100 µL full melanoma medium with 4 µM Caspase-3/7 activity dye (CellEvent, Thermo Fisher Scientific). Plates were then imaged on a Celigo Imaging Cytometer (Nexcelom) for a t0 count of viable target cells. Two days prior to coculture, TILs were thawed into complete TIL media with 3000 IU/mL IL-2. TILs were collected and resuspended in complete melanoma medium and added to melanoma cells in 100 µL per well at increasing TIL:target ratios. After 24 h, 48 h, and 72 h, plates were imaged again to track viable melanoma cell growth. At the end of co-culture, plates were spun at 400 x g for 5 min and cleared culture media was collected for cytokine analysis by ELISA described below. Cell counts were performed using Celigo Imaging Cytometer software. Viable melanoma cells were defined as DsRed+ cells that were not Caspase-3/7+, with a size threshold to exclude debris. Viable melanoma cell counts were then normalized to respective t0 counts. Where indicated, target cells were pretreated overnight with 10 µg/mL anti-CD58 TS2/9 blocking antibody or 10 µg/mL anti-PD-L1 (29E.2A3) blocking antibody or IgG isotype control, and treated again upon addition of T cells. T cells were pretreated with 200 ng/mL anti-CD2 LoCD2b blocking antibody or IgG isotype control overnight and then again upon addition to target cells, where indicated. For PD-L1 blocking co-culture experiments, TILs were cultured overnight on plates coated with 1 µg/mL anti-CD3 (OKT3) activating antibody prior to addition to target cells. TILs were then assayed for PD-1 expression by flow cytometry, as described below.

### Flow Cytometry

For analysis of cell surface markers, cells were detached with Accutase (Innovative Cell Technologies, Inc.) as needed and washed with PBS. Cells were first stained with Zombie-NIR Fixable Viability Dye (Biolegend) for 15 min at RT in the dark and washed once with PBS, followed by staining with fluorophore-conjugated antibodies in FACS buffer (2% FBS, 2 mM EDTA in PBS) for 20 min on ice in the dark. Cells were washed 2X with FACS buffer and then fixed with Fixation Buffer (Biolegend), washed 2X with FACs buffer, and analyzed on an Aurora 3-laser spectral cytometer (Cytek). Cells stained with isotype control antibodies were used as negative controls. All analysis of flow cytometry data was performed using FlowJo (BD).

### SDS-PAGE and immunoblotting

For each blot, melanoma cells were stimulated with or without 10 ng/mL IFN-ɣ for 72 h and detached and collected using 0.05% Trypsin (Thermo). Cells were lysed in RIPA buffer (Sigma) with 10X concentration of Halt™ Protease and Phosphatase Inhibitor Cocktail (Thermo) for 30 min on ice, and then spun at 10,000 rpm for 15 min at 4C in a microcentrifuge. Supernatants were collected and protein was quantified using the Pierce™ BCA Protein Assay Kit (Thermo). Cell lysates brought to the desired concentration in RIPA buffer with 1X Laemmli SDS-Sample Buffer (Boston Bioproducts Inc.) and boiled at 95 ºC for 10 min. Lysates were ran on 10% TGX™ FastCast acrylamide gels (Bio-Rad) in tris-glycine SDS running buffer and transferred onto PVDF membranes using the iBlot 2 Dry Blotting system (Thermo). Blots were blocked with 1X Western Blocking Reagent (Sigma) in tris-buffered saline (TBS) for 45 min and then probed overnight with primary antibodies diluted in TBS with 0.1% Tween-20 (TBS-T; Sigma) at 4 ºC with rotation. Blots were then washed 3X with TBS-T and probed with fluorophore-conjugated secondary antibodies (Li-Cor) for 1 h at RT with rocking and washed 3X with TBS-T. All blots were developed on the Odyssey CLx imaging system (Li-Cor) and analyzed in Image Studio Lite (Li-Cor).

### T cell co-stimulation proliferation assay

24-well tissue culture-treated plates were pre-coated with 1 µg/mL anti-CD3 (OKT3) antibody (Miltenyi Biotec 130-093-387) in 1 mL PBS per well, +/-1 µg/mL anti-CD28 (CD28.2) antibody (eBioscience/Thermo 16-0289-81) or recombinant human CD58-Fc chimera protein (R&D Systems 10068-CD-050) at 37 ºC for 2 hours. TILs were stained with 1 µM CFSE (Thermo C34570) in PBS for 1 min at 37 ºC at a cell density of 5 million cells/mL and then washed with complete media. Cells were then seeded at a density of 300,000 cells/well in 1 mL TIL media. Cleared cell media was collected after 24 and 48 h to use for ELISAs. Five days after seeding, TILs were collected, stained with Zombie-NIR (Biolegend), fixed with Fixation Buffer (Biolegend), and analyzed for cell proliferation by flow cytometry.

### ELISAs

IL-2 and IFN-ɣ levels were detected in cleared cell media using the ELISA MAX™ Deluxe ELISA kits (Biologend) per manufacturer protocol. Absorbance values were measured using a Synergy H1 plate reader (Biotek).

### In vivo melanoma xenograft, tumor processing, and analysis

32 female NOD.Cg-*Prkdc*^*scid*^ *Il2rg*^*tm1Sug*^ Tg(CMV-IL2)4-2Jic/JicTac mice (Taconic) were implanted with 2 million each MaMel134 DsRed control and MaMel134 DsRed *CD58* KO (n=16) or *CD58*-TM OE (n=16) cells resuspended in 25% Matrigel (Corning) via bilateral subcutaneous flank injections. Mice were weighed once a week and tumors were measured 2-3 times/week using digital calipers. Three days prior to ACT/PBS injection, cryopreserved TILs were thawed and cultured in T cell media supplemented with 300 IU/ml human recombinant IL-2. Once tumors were palpable for all mice (∼1 month), 8 mice from each cohort were injected with 5 million MaMel134 TILs in 100 µL PBS by tail vein injection, with the remaining half of each cohort receiving 100 µL PBS control. Submandibular blood collection was performed once a week to monitor circulating TILs by flow cytometry. Due to unclear responses to ACT, one month after initial ACT/PBS injections, a second round of ACT or PBS was administered. Once largest tumor pairs reached a summed diameter of 20 mm, mice were sacrificed, and tumors were harvested. Tumors were weighed and dissociated with digestion media (collagenase D and DNase I (both Roche, Sigma-Aldrich) in PBS) and filtered through 70 µM cell strainers to produce a single cell suspension for flow analysis. Cells were incubated in ACK Lysing Buffer (Quality Biological) for 1 min at RT and washed with PBS; ACK lysis was repeated as necessary. Cells were stained with Zombie-NIR followed by Fc block with anti-mouse CD16/32 antibody (Biolegend). Cells were then stained for cell surface markers as described above. Additional intracellular markers were stained using the Foxp3/Transcription Factor Staining Kit (eBioscience, Thermo) per manufacturer protocol. FMO controls were prepared for Ki-67 and CD28. Tumor volumes were calculated as (Tumor width x Tumor width x Tumor length). Fold change growth was calculated using the average of the three tumor measurements immediately before, after, and on the day of the first ACT/PBS injection as t0.

### A375 Cas9 transduction

pLX_311-Cas9 lentivirus was produced via transfection of HEK293T cells as described above (plasmid gift of William Hahn and David Root; Addgene #118018). A375 WT cells were seeded in 12-well plates with 1 million cells per well with increasing volumes of pLX_311-Cas9 lentivirus to a final volume of 2 mL with 4 µg/mL polybrene. Cells were spun at 1,000 x g for 2 h at 30 ºC, and 2 mL media was added to each well after spinning. Cells were then incubated at 37 ºC, 5% CO_2_ in a humidified incubator overnight. The next morning, 100,000 transduced cells and non-transduced cells were seeded at equal densities across two wells each per condition of a 6-well plate. The next day, cells were put into selection with 4 µg/mL blasticidin (Thermo), and at the end of selection transduction efficiency was calculated as above.

### Cas9 activity assay

To measure Cas9 editing efficiency, Cas9-expressing A375 cells were transduced with lentivirus containing expression plasmid pXPR-011 (gift of John Doench and David Root; Addgene #59702), which encodes for enhanced green fluorescent protein (EGFP) and a guide RNA targeting eGFP^48^. As a no-editing control, WT A375 cells were also transduced. Transduced cells were put into 0.75 µg/mL puromycin selection. At the end of selection, EGFP expression of transduced cells was assessed by flow cytometry on an Aurora spectral cytometer (Cytek). Cas9 activity was measured as the percentage of EGFP-negative cells, using no-editing control cells as gating controls.

### Brunello library titration

A375 Cas9-expressing cells were seeded in 12-well plates at a density of 1 million cells per well with increasing volumes of lentivirus (1.02×10^7^ transduction units/mL) carrying the human CRISPR KO pooled library Brunello in a lentiGuide-Puro backbone (gift from David Root and John Doench; Addgene #73178) to a final volume of 2 mL with 4 µg/mL polybrene. Cells were spun at 1,000 x g for 2 h at 30 ºC, and 2 mL media was added to each well after spinning, keeping cells in 4 µg/mL blasticidin selection. Cells were then incubated at 37 ºC, 5% CO2 in a humidified incubator overnight. The next morning, 200,000 transduced cells and non-transduced cells were seeded across two wells each per condition of a 6-well plate. The next day, cells were put into selection with 0.75 µg/mL puromycin, and at the end of selection transduction efficiency was calculated as above. Cells with a transduction efficiency of ∼20% were selected, resulting in an MOI of 0.3 and therefore a single integration rate of 95%.

### Anti-CD58-APC antibody titration

The anti-CD58-APC antibody used for the CRISPR-Cas9 KO screen was titrated by staining pooled A375 WT and *CD58* KO cells as above with Zombie-Violet followed by increasing concentrations of anti-CD58-APC (Biolegend) antibody. Cells were then fixed and analyzed on an Aurora spectral cytometry (Cytek). For each condition, CD58-negative and CD58-positive populations were gated the median fluorescence intensity of APC fluorescence was calculated. The separation index for each antibody concentration was then calculated as follows:

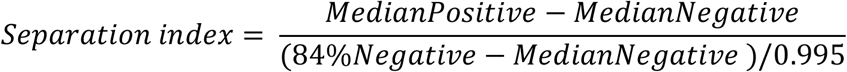

The lowest concentration that gave the relatively highest separation index was chosen and used for the screen.

### Transduction of Brunello library

To reach a desired representation of ∼500 cells/sgRNA, with a transduction efficiency of 20%, 220 million A375 Cas9-expressing cells were seeded across 12-well plates at a density of 1 million cells/mL with 30 µL Brunello virus per well to a final volume of 2 mL with 4 µg/mL polybrene. Cells were spun at 1,000 x g for 2 h at 30 ºC. After spinning, 2 mL fresh media was added to each well and cells were incubated overnight at 37 ºC, 5% CO2 in a humidified incubator. One day after transduction, cells were expanded up into T175 flasks. At two days post-transduction, cells were put into selection with 0.75 µg/mL puromycin, and cells were continuously expanded up until sorting, keeping in puromycin and blasticidin throughout to maintain Cas9 and sgRNA expression. Two biological replicates were performed.

### A375 Cas9 Brunello CD58 FACS

10 days after transduction with Brunello virus, sgRNA library- and Cas9-expressing A375 cells were detached with Accutase, washed with PBS, and split into two technical replicates. For each replicate, 130 million cells were transferred to 96 well V-bottom storage plates (Corning) with 1 million cells/well. Cells were first stained with Zombie-Violet (Biolegend) for 15 min at RT and washed 1X. Cells were then stained with anti-CD58-APC antibody in FACS buffer for 20 min on ice and washed 2X. Cells were fixed with fixation buffer and incubated for 20 min at RT and washed 2X. Cells were then resuspended in 100 µL FACS buffer/well and pooled together to use for sorting. Cells were sorted on an Influx (BD) cell sorter. A375 Cas9 cells without sgRNA expression and A375 *CD58* KO cells were used for gating controls. Cells with the lowest 5% of CD58 expression were collected as the target population, alongside cells within ∼1 standard deviation above and below the median as controls. Sorted cells were kept in FACS buffer and stored at 4 ºC overnight. Unsorted cells were also reserved to be used for essential gene dropout analysis.

### A375 Cas9 Brunello genomic DNA (gDNA) extraction

gDNA was collected from sorted and unsorted cells using a DNeasy Blood and Tissue Kit (Qiagen). Cells were first pelleted and split across multiple microcentrifuge tubes with a maximum of 2.5 million cells/tube. Cells were resuspended in 200 µL PBS, to which 20 µL proteinase K and 200 µL Buffer AL were added. Cells were then incubated overnight at 65 ºC with shaking at 1000 rpm to remove formaldehyde crosslinking. The next morning, gDNA was extracted from cell lysates per manufacturer protocol.

### sgRNA PCR amplification and clean up

sgRNA sequences were amplified from gDNA by PCR using Ex Taq DNA polymerase (TaKaRa). PCR mixes were prepared with 10 µL 10X reaction buffer, 8 µL dNTP, 5 µL DMSO, 0.5 µL P5 primer mix (100 µM), 1.5 µL Ex Taq polymerase, 5 µL unique P7 primer (10 µM) per condition, and a maximum of 10 µg gDNA to a final volume of 100 µL. PCR reactions were cycled at 95 ºC for 1 minute; followed by a total 28 cycles of 95 ºC for 30 sec, 53 ºC for 30 sec, and 72 ºC for 30 sec; 72 ºC for 10 minutes; and then finally 4 ºC. 25 µL from each PCR product was pooled and loaded onto a 1.5% agarose gel (Bio-Rad) and ran at 100V. The ∼350 bp PCR product was extracted using the Zymoclean Gel DNA Recovery Kit (Zymo) per manufacturer protocol. To remove all buffer salts, a final inclusive cleanup was performed using SPRI beads (Agilent). SPRI beads were added to PCR product at a 1.2X ratio and incubated for 5 min. After placing on a magnet for two minutes, beads were washed 2X with 80% ethanol and resuspended in TE buffer and quantified on a TapeStation (Agilent).

### CRISPR/Cas9 KO screen sequencing and analysis

Final prepared sgRNA library was sequenced on a NextSeq 500 sequencer (Illumina). Data was analyzed using MAGeCK Version 0.5.9.2. Briefly, sgRNA counts by condition were generated using mageck count and compared with mageck test using default parameters. Comparisons between FACS gates identified potential positive regulators of CD58. Inclusion criterium for defined positive regulators of CD58 was <0.25 FDR for enrichment of a gene target in CD58^lo^ v. CD58^mi^ cells across all four replicates of the screen.

### Co-immunoprecipitation

For each IP, 5 million melanoma cells were stimulated with or without 10 ng/ml IFN-ɣ for 72 h and detached using 0.05% Trypsin and pelleted. Cells were then lysed in 1% digitonin lysis buffer (50 mM Tris-HCl, pH 8.0, 150 mM NaCl, 5 mM EDTA, 1% digitonin, 10X Halt™ Protease and Phosphatase Inhibitor Cocktail (Thermo)) for 30 min at 4 ºC with rotation and then spun at 13,000 x g for 10 min. Supernatant was collected and lysates were then diluted 2X to 0.5% digitonin. Lysates were incubated with 2 µg anti-CD58 or anti-CMTM6 antibody for 3 h at 4 ºC with rotation. Antigen-antibody mixtures were then used to resuspend 10 µL cleared Protein A or Protein G Dynabeads (Thermo) and incubated for 2 h at 4 ºC with rotation. Samples were washed 4 times on a DynaMag™-2 (Thermo) with wash buffer (50 mM Tris-HCl pH 8.0, 150 mM NaCl, 5 mM EDTA, 0.1% digitonin). Samples were then either flash frozen on dry ice for MS analysis, or resuspended in 1X Laemmli SDS-Sample Buffer (Boston BioProducts) and boiled at 95 ºC for 10 min to elute off protein for SDS-PAGE as described above. For MS analysis, three replicates were independently prepared of CD58 and IgG control pulldowns.

### On-bead digestion of samples for mass spectrometry

Proteins bound to magnetic beads were washed five times with 200 µl of 50 mM ammonium bicarbonate and subjected to disulfide bond reduction with 5 mM TECP (RT, 30 min) and alkylation with 10 mM iodoacetamide (RT, 30 min in the dark). Excess iodoacetamide was quenched with 5 mM DTT (RT, 15 min). Proteins bound on beads were digested overnight at 37°C with 1 µg of trypsin/LysC mix. The next day, digested peptides were collected in a new microfuge tube and digestion was stopped by the addition of 1% TFA (final v/v), and centrifuged at 14,000 g for 10 min at room temperature. Cleared digested peptides were desalted on SDB-RP Stage-Tip and dried in a speed-vac. Peptides were dissolved in 3% acetonitrile/0.1% formic acid.

### Liquid chromatography with tandem mass spectrometry (LC-MS/MS)

Peptides were separated within 80 min at a flow rate of 400 nl/min on a reversed-phase C18 column with an integrated CaptiveSpray Emitter (25 cm x 75µm, 1.6 µm, IonOpticks). Mobile phases A and B were with 0.1% formic acid in water and 0.1% formic acid in ACN. The fraction of B was linearly increased from 2 to 23% within 70 min, followed by an increase to 35% within 10 min and a further increase to 80% before re-equilibration. The timsTOF Pro was operated in PASEF mode^49^ with the following settings: Mass Range 100 to 1700m/z, 1/K0 Start 0.6 V·s/cm2, End 1.6 V·s/cm2, Ramp time 100ms, Lock Duty Cycle to 100%, Capillary Voltage 1600V, Dry Gas 3 l/min, Dry Temp 200°C, PASEF settings: 10 MSMS Frames (1.16 seconds duty cycle), charge range 0-5, active exclusion for 0.4 min, Target intensity 20000, Intensity threshold 2500, CID collision energy 59 eV. A polygon filter was applied to the *m/z* and ion mobility plane to select features most likely representing peptide precursors rather than singly charged background ions.

### LC-MS/MS data analysis

Acquired PASEF raw files were analyzed using the MaxQuant environment v.2.0.1.0 and Andromeda for database searches at default settings with a few modifications^50^. The default is used for first search tolerance and main search tolerance (20 ppm and 4.5 ppm, respectively). MaxQuant was set up to search with the reference human proteome database downloaded from UniProt. MaxQuant performed the search trypsin digestion with up to 2 missed cleavages. Peptide, site, and protein false discovery rates (FDR) were all set to 1% with a minimum of 1 peptide needed for identification; label free quantitation (LFQ) was performed with a minimum ratio count of 1. The following modifications were used for protein identification and quantification: Carbamidomethylation of cysteine residues (+57.021 Da) was set as static modifications, while the oxidation of methionine residues (+15.995 Da), deamidation (+0.984) on asparagine and glutamine were set as a variable modification. Results obtained from MaxQuant, were imported into Perseus v.1.6.15.0^51^ for t-test statistical analysis (FDR<0.05) to identify proteins demonstrating statistically significant changes in abundance. Inclusion criteria for selection as an interactor with CD58: FDR<0.05, log2(foldchange) of LFQ intensity value >3, average spectral count within CRAPome database <2 to exclude background contaminants^52^.

### RT-qPCR

Melanoma cells were stimulated with or without 10 ng/mL IFN-ɣ for 72 h and then detached using 0.05% Trypsin and pelleted. Total RNA was collected from cell pellets using an RNeasy Mini kit (Qiagen). cDNA was then generated using the iScript™ cDNA Synthesis Kit (BioRad), and qPCR was performed using TaqMan Fast Advanced Master Mix (Applied Biosystems) in combination with TaqMan Gene Expression Assays per manufacturer protocol on a ViiA 7 RT-PCR system (Applied Biosystems). The assay IDs for TaqMan Gene Expression Assays used were as follows: Hs00204257_m1 (*CD274*), Hs01560660_m1 (*CD58*), Hs99999903_m1 (*ACTB*), and Hs02786624 (*GAPDH*). Gene expression was normalized to *ACTB* and *GAPDH* expression, and compared to WT, unstimulated cells.

### Degradation Assay

2686 melanoma cells were assayed for degradation of CD58, PD-L1, and HLA-A,B,C as previously described^30^. Briefly, cells were cultured for 72 h with or without 10 ng/mL IFN-ɣ and then stained with anti-CD58, anti-PD-L1, and anti-HLA-A,B,C conjugated antibodies for 1 h on ice. After washing, cells were replated in complete medium and incubated at 37 ºC for 3-6 h in the presence or absence of 50 µM chloroquine or 50 nM concanamycin A. Cells were collected at each timepoint with Accutase, washed, stained with Zombie-NIR, fixed as described previously, and analyzed on an Aurora spectral cytometer (Cytek). Degradation of CD58, PD-L1, and HLA-A,B,C was measured by a decline in fluorescence intensity over time.

### Immunocytochemistry

2686 melanoma cells were seeded in 8-well chambered coverglass slides (CellVis) at a density of 20,000 cells per chamber with or without 10 ng/mL IFN-ɣ and incubated overnight at 37 ºC, 5% CO2 in a humidified incubator. The following day, cells were fixed in 4% paraformaldehyde (Fisher Scientific) for 15 min at RT. Cells were washed 2X with PBS and permeabilized with 0.25% Triton X-100 (Fisher Scientific) for 5 min at RT. Cells were washed 2X and then blocked with 1% BSA (Corning) for 1 h at RT. Cells were washed 2X and treated with freshly made 1 mg/mL NaBH4 in PBS for 7 min at RT. Cells were washed 3X and then stained with primary antibody prepared in 1% BSA for 1 h at RT. Cells were washed 3X and then stained with secondary antibody for 30 min at RT. Cells were washed 3X and then optionally stained with fluorophore-conjugated antibody for 1 h at RT. Cells were washed 3X and stained with Hoechst 33342 (Thermo) for 5 min at RT. Cells were washed and then imaged using an LSM 900 confocal microscope (Zeiss). All image analysis was performed in ImageJ^53^.

### Antibodies

Antibodies used listed below.

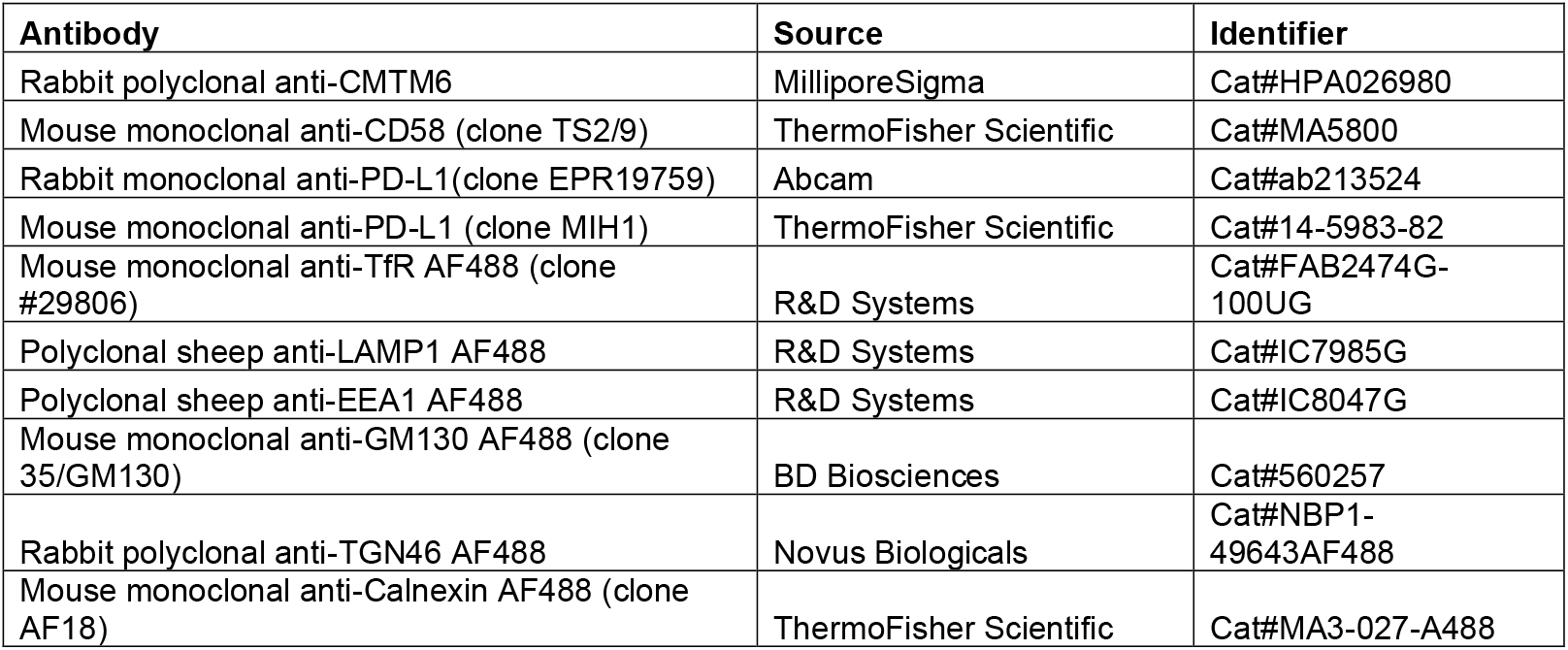
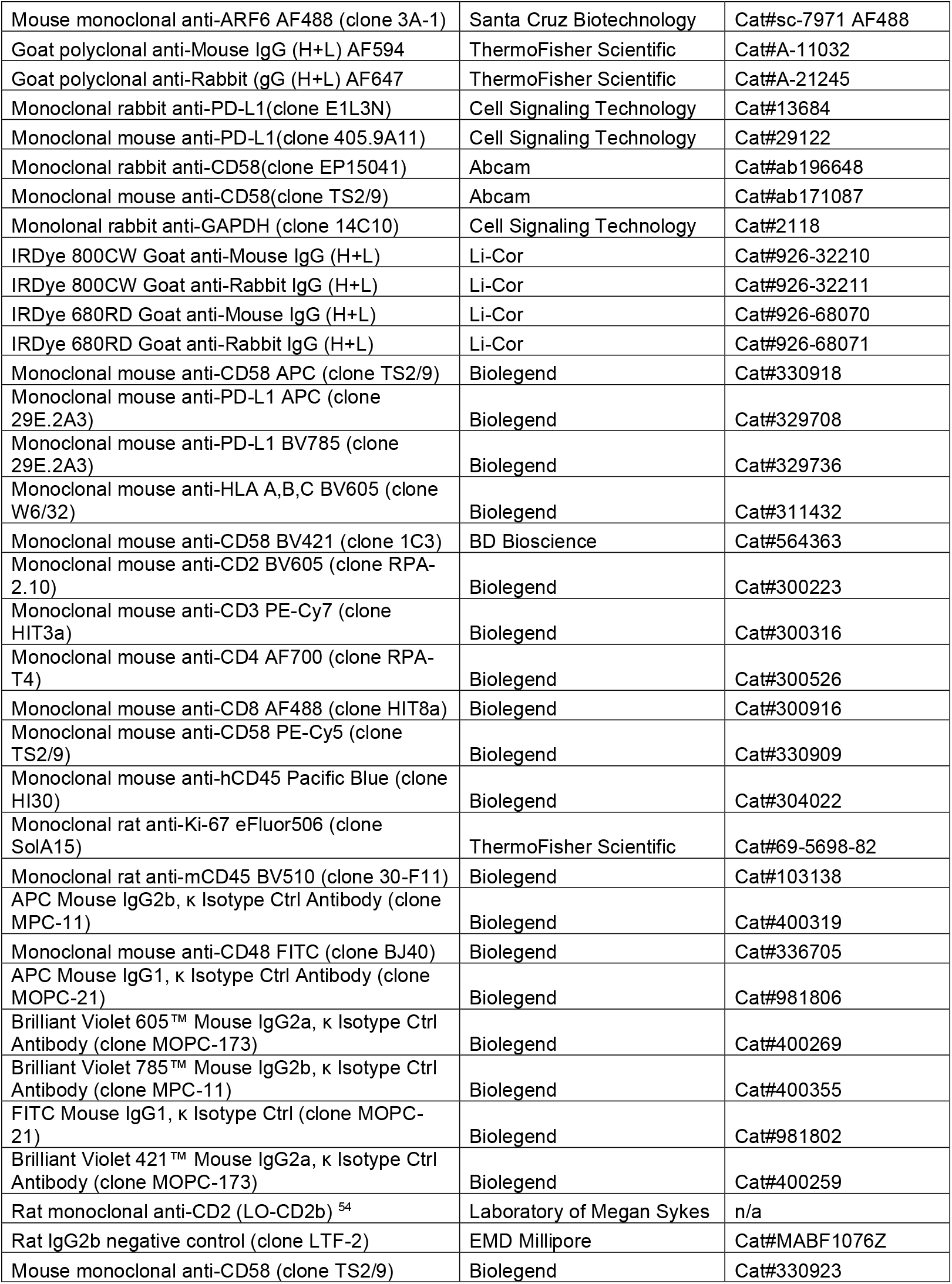
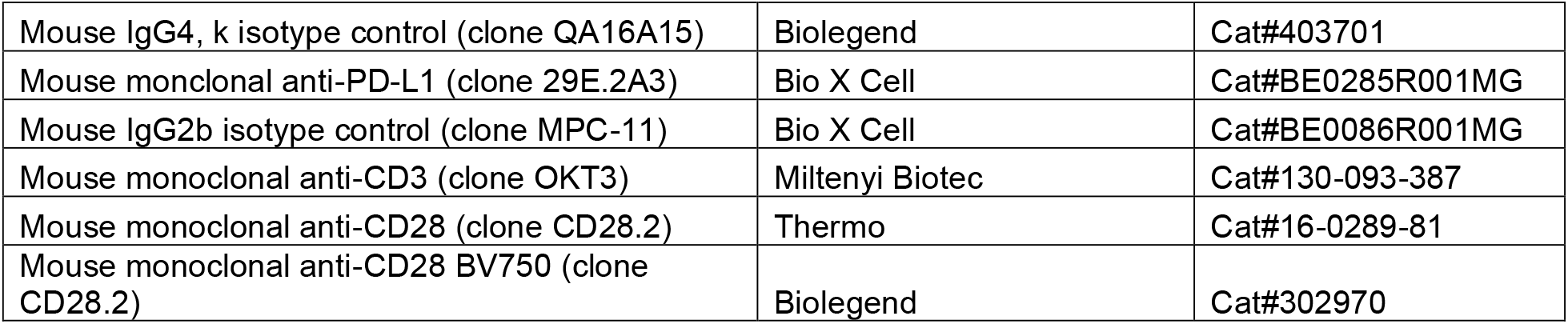

