## Supplemental Figures for "The CD58:CD2 axis is co-regulated with PD-L1 via CMTM6 and governs anti-tumor immunity"

### Figure S1: RNA-seq analysis of patient tumors across cancer types

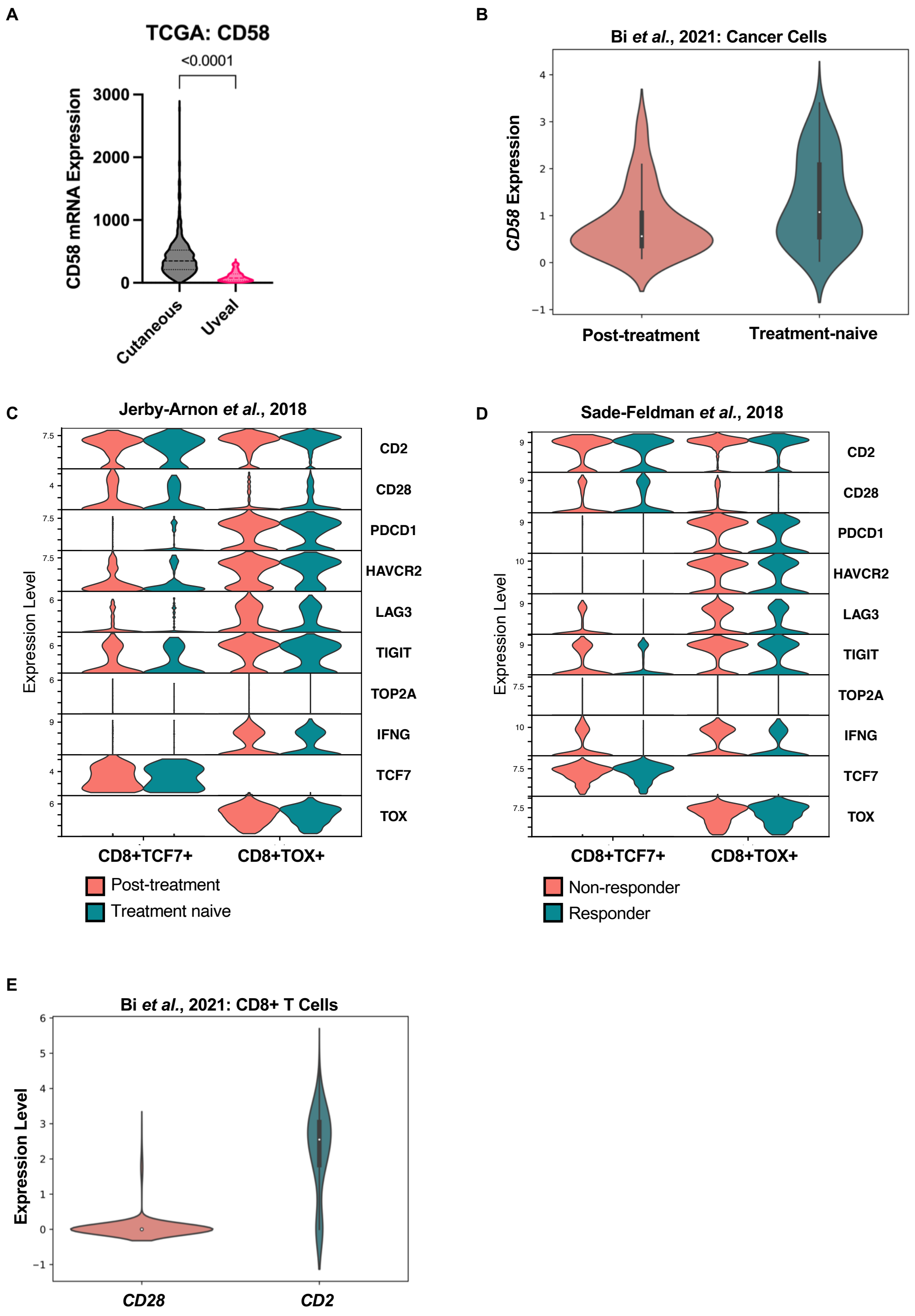

**Figure S2: CD58-CD2 ligation is critical to T-cell-mediated killing across cell lines**

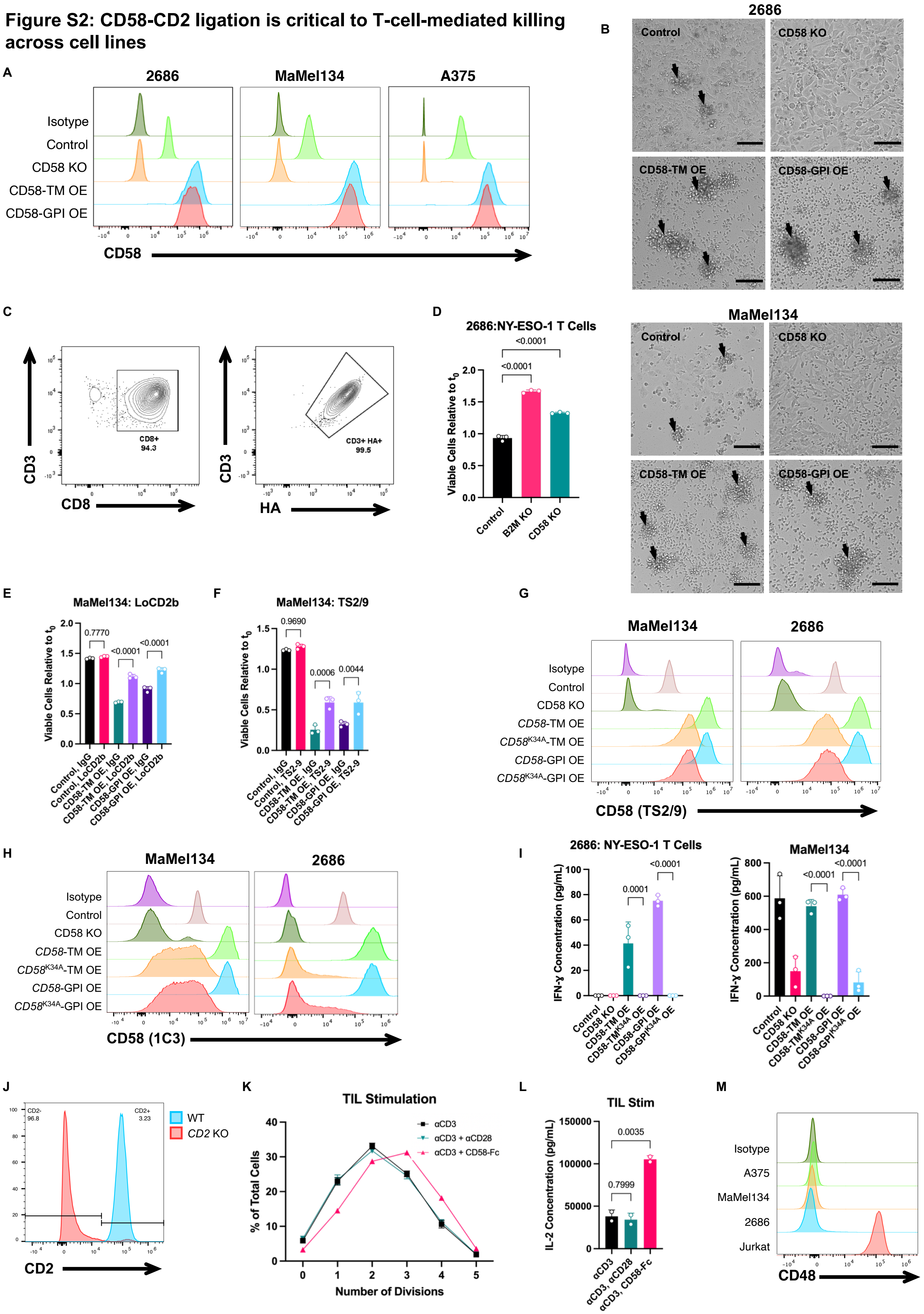

**Figure S3: *In vivo* model of ACT-treated melanoma recruits intratumoral CD8+ TILs**

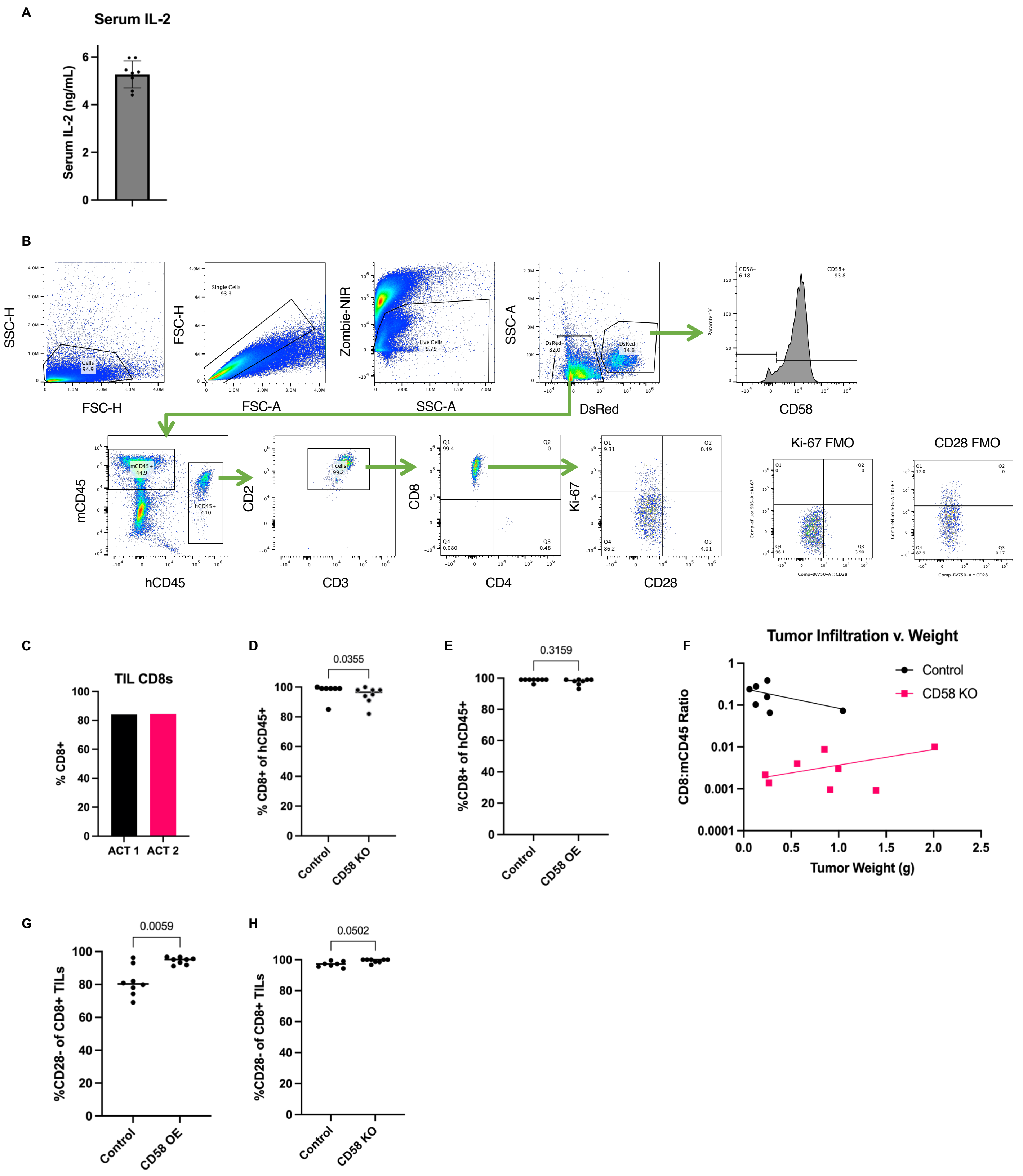

#### Figure S4: CD58 regulates PD-L1 expression across multiple cell lines

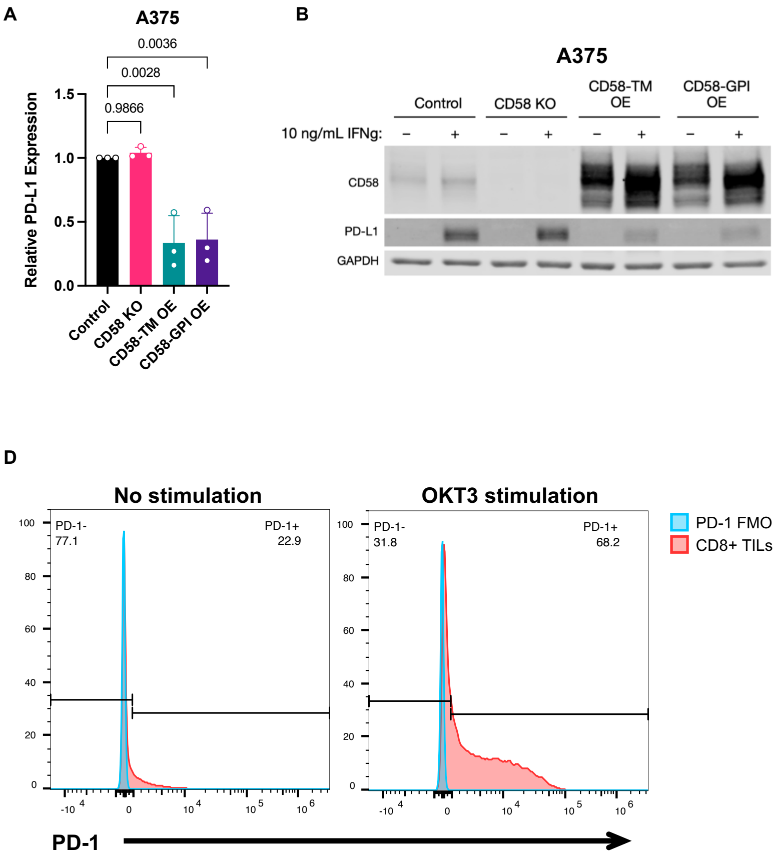

### Figure S5: Preparation and analysis of CRISPR/Cas9 KO screen to identify regulators of CD58

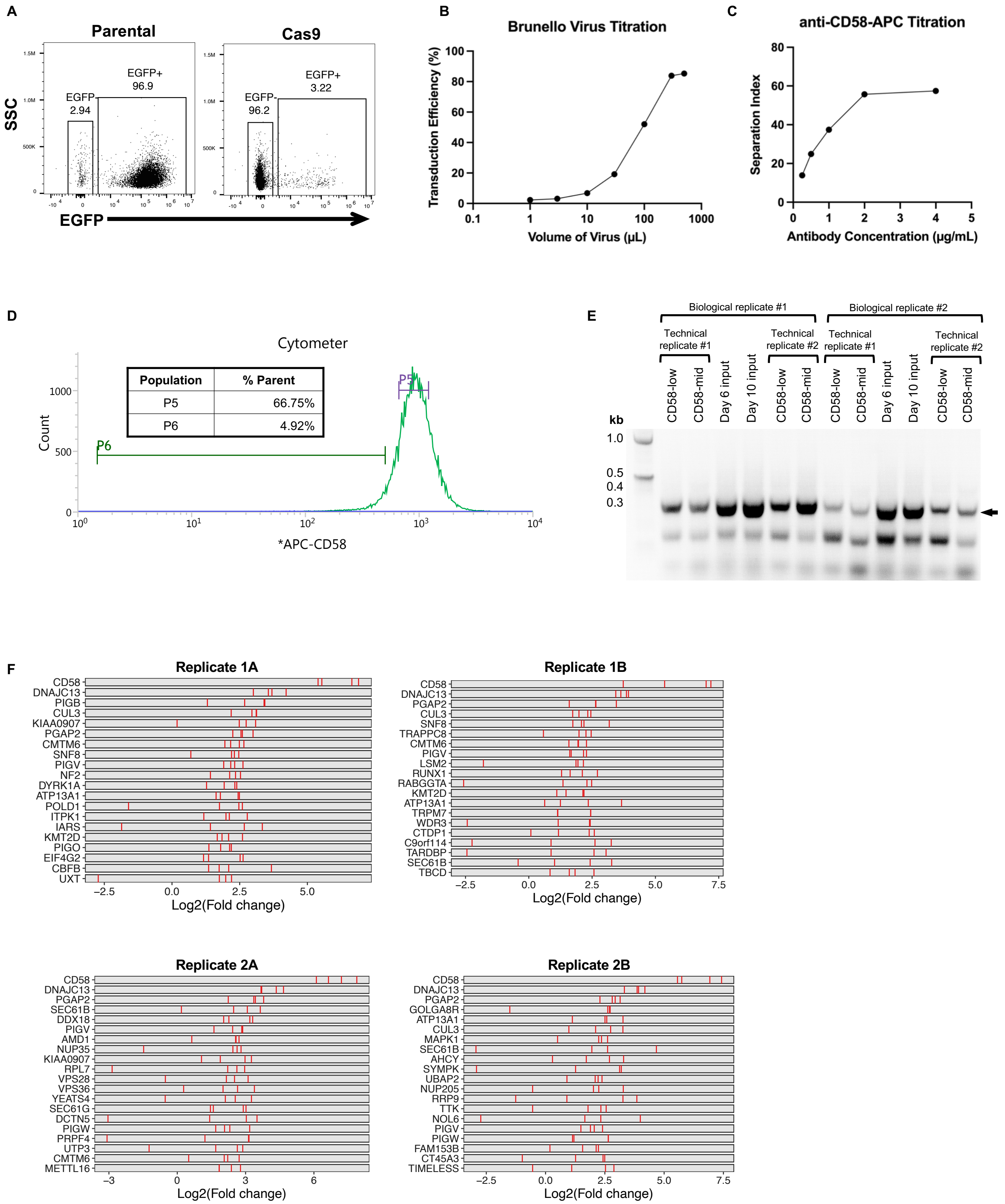

### Figure S6: CMTM6 regulates CD58 at a protein level across multiple cell lines

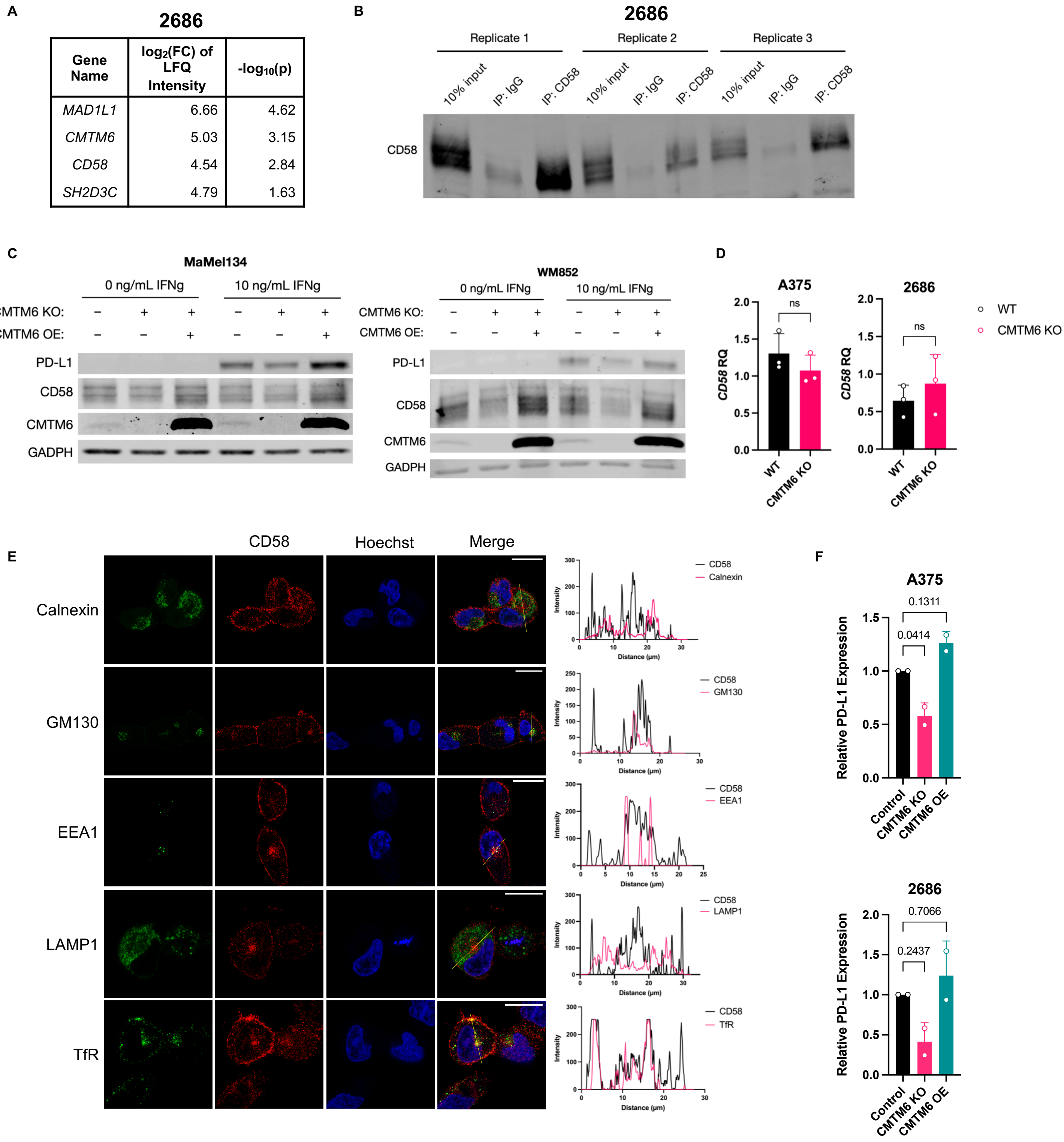

#### Figure S7: CMTM6 is critical for CD58's regulation of PD-L1

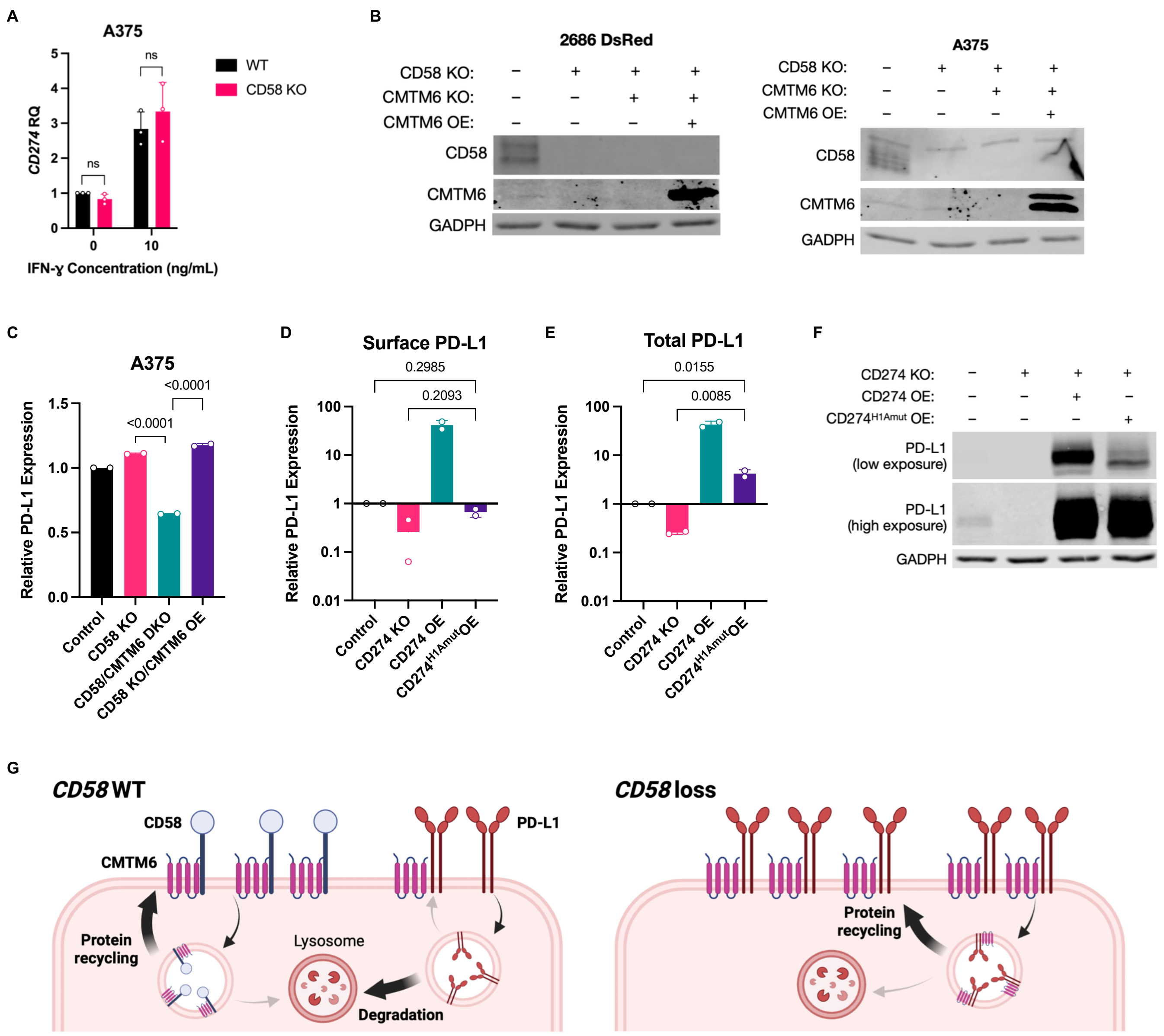
